# Collateral sensitivity as a strategy to suppress resistance emergence: the challenge of diverse evolutionary pathways

**DOI:** 10.1101/2022.04.07.486819

**Authors:** Rebecca E.K. Mandt, Madeline R. Luth, Mark A. Tye, Ralph Mazitschek, Sabine Ottilie, Elizabeth A. Winzeler, Maria Jose Lafuente-Monasterio, Francisco Javier Gamo, Dyann F. Wirth, Amanda K. Lukens

## Abstract

Drug resistance remains a major obstacle to malaria control and eradication efforts, necessitating the development of novel therapeutic strategies to treat this disease. Drug combinations based on collateral sensitivity, wherein resistance to one drug causes increased sensitivity to the partner drug, have been proposed as an evolutionary strategy to suppress the emergence of resistance in pathogen populations. In this study, we explore collateral sensitivity between compounds targeting the *Plasmodium* dihydroorotate dehydrogenase (DHODH). We profiled the cross-resistance and collateral sensitivity phenotypes of several DHODH mutant lines to a diverse panel of DHODH inhibitors. We focus on one compound, TCMDC-125334, which was active against all mutant lines tested, including the DHODH C276Y line, which arose in selections with the clinical candidate DSM265. We found that selection of the DHODH C276Y mutant with TCMDC-125334 yielded additional genetic changes in the *dhodh* locus. These double mutant parasites exhibited decreased sensitivity to TCMDC-125334 and were highly resistant to DSM265. Finally, we tested whether collateral sensitivity could be exploited to suppress the emergence of resistance in the context of combination treatment by exposing wildtype parasites to both DSM265 and TCMDC-125334 simultaneously. This selected for parasites with a DHODH V532A mutation which were cross-resistant to both compounds and were as fit as the wildtype parent *in vitro*. The emergence of these cross-resistant, evolutionarily fit parasites highlights the mutational flexibility of the DHODH enzyme.

## Introduction

Antimicrobial resistance threatens our ability to combat multiple infectious agents, and is widely recognized as one of the greatest public health threats of the 21^st^ century ^1^. Globally, drug resistance is a recurring obstacle in efforts to control many endemic infectious diseases, and malaria is no exception. The emergence of resistance in *Plasmodium* parasite populations and subsequent treatment failure has been documented for all antimalarial drugs in clinical use, including frontline artemisinin combination therapies (ACTs) ^2^. Thus, there is an ongoing need to develop a next generation of drugs targeting distinct vulnerabilities of *Plasmodium* parasites.

Additionally, understanding the pathways and susceptibility to drug resistance is important to assess next-generation antimalarials, ideally while they are still early in the development process. Inhibitors targeting the *Plasmodium* dihydroorotate dehydrogenase (DHODH) offer an illustrative example. DHODH is an electron transport chain protein that catalyzes the rate-limiting step of pyrimidine biosynthesis and is essential for parasite growth and survival ^3^. While the clinical DHODH inhibitor candidate DSM265 showed promising activity in pre-clinical and clinical studies^4-8^, we previously demonstrated that resistance to DSM265 emerges rapidly both *in vitro*, and in a humanized mouse model of *P. falciparum* infection^9^. Resistance was primarily conferred by point mutations in the *dhodh* locus. In Phase 2 clinical trials with DSM265, two out of 24 patients failed treatment due to resistance, with recrudescent parasites harboring mutations in DHODH including DHODH C276Y, C276F and G181S ^4^. The C276Y and C276F mutations were also observed in *in vitro* resistance selections, as was DHODH G181C^9-11^. The point mutations C276F and G181D were observed in our *in vivo* selection model^9^.

A common strategy to counter the evolution of resistance is to utilize a combination of two or more drugs with distinct antimicrobial mechanisms. The basic rationale for this strategy is that even if there is some probability of an organism acquiring resistance to each drug separately, the probability that two resistance mutations would occur simultaneously is much lower. Combination therapy is the standard of treatment for malaria ^12^, tuberculosis ^13^, and HIV/AIDS^14^, and is increasingly being recommended for the treatment of other bacterial infections ^15-17^. However, multi-drug resistance has emerged in many major human pathogens, complicating patient treatment and threatening public health efforts to control the spread of disease ^18-21^. Large-scale bacterial evolution studies have also shown that certain combinations of drugs can actually accelerate the emergence of resistance compared to single drug treatment ^22,23^. Thus, there is a need to re-evaluate the current treatment approach, and to strategically design combination therapies based on their ability to slow or suppress the emergence of resistance.

One possible strategy takes advantage of collateral sensitivity, in which resistance to one drug causes increased sensitivity to another ^24^. The phenomenon of collateral sensitivity, or ‘negative cross-resistance,’ has been observed across various antibiotic classes in bacteria, as well as anti-fungals, and even cancer therapies ^25-31^. Laboratory studies evaluating the evolution of resistance to antibiotics have demonstrated that when resistance to one drug leads to collateral sensitivity to another, the two drugs can be used in a sequential cycling strategy to maintain sensitive bacterial populations ^25,32^, or simultaneously in combination to slow or suppress the evolution of resistance ^33-35^.

Collateral sensitivity to various classes of drugs targeting *Plasmodium* parasites has also been observed. For example, isoforms of the chloroquine resistance transporter (*Pf*CRT) that transport chloroquine out of the parasite’s digestive vacuole can cause increased sensitivity to a variety of compounds ^36-41^. Collateral sensitivity has also been observed between compounds targeting different subunits of the *Plasmodium* proteasome *Pf*20S ^42,43^, between different chemotypes targeting the P-Type Cation-Transporter ATPase 4 (*Pf*ATP4) ^44^, and between compounds targeting the Q_o_ and Q_i_ sites of cytochrome b ^45^. We have also previously shown that collateral sensitivity occurs during the evolution of resistance to *Plasmodium* DHODH inhibitors. Mutations in DHODH conferring resistance to one inhibitor can alter the enzyme such that it becomes more sensitive to inhibition by other, structurally-distinct compounds ^36,46^. The existence of collateral sensitivity among a wide range of antimalarial inhibitors suggests a promising opportunity to exploit this phenomenon to block the emergence of resistant parasites ^36^.

In an effort to expand this work, we screened antimalarial libraries from GSK, and successfully identified several compounds with increased activities against DHODH mutants relative to wildtype parasites ^47^. In this study, we test the hypothesis that combining two DHODH inhibitors based on their collateral sensitivity profiles would suppress the emergence of resistant parasites. We focused on the compound TCMDC-125334, a DHODH inhibitor that exhibited increased activity against all mutant lines tested, including those that arose during selection with the clinical candidate DSM265. We used *in vitro* resistance selections to characterize the pathways to resistance for TCMDC-125334 and then tested whether parasites develop resistance in the context of combined treatment with DSM265 and TCMDC-125334.

## Results

### TCMDC-125334 demonstrates activity against all DHODH mutant parasite lines tested

Previously, we screened GSK chemical libraries from the Tres Cantos Antimalarial Set for activity against either wildtype or DHODH mutant parasites (Table S1) ^47^. In an extended cross-resistance analysis, we characterized the activity of 17 compounds against eight DHODH mutant lines (Figure 1A, Table S1, Figure S1, Data Table S1). We visualized the log-transformed fold-change in EC_50_ of each mutant relative to its parental line on a heatmap (Figure 1B, Figure S1, Data File S1). Hierarchical clustering of this data reveals broad patterns of cross-resistance and collateral sensitivity. The DHODH C276Y, F227I, F227I/L527I, F227L, and F227L/L31F mutant lines have similar dose response phenotypes. The DHODH E182D and I263F mutant lines also share similar cross-resistance profiles, as we previously reported ^47^. In contrast, the DHODH L531F mutant line has a unique sensitivity phenotype (Figure 1B). This analysis also suggests some structure-activity relationships. Five related triazolopyrimidine-based compounds (DSM265, DSM267, TCMDC-124402, TCMDC-124417, TCMDC-123826) cluster together, as all mutant lines show cross-resistance against these molecules (Figure 1B). TCMDC-123823, TCMDC-123545, and TCMDC-123566 also cluster together based on their activity profile, and have a similar chemical structure (Figure 1B, Data Table S1). Based on our analysis, there was one compound, TCMDC-125334, that was active against all parasite lines tested. The eight mutant lines had varying degrees of sensitivity to this compound, from DHODH I263F which is 1.3-fold more sensitive than the wildtype parent, to DHODH C276Y, which is 12-fold sensitive. In contrast, all mutant lines were at least 10-fold resistant to DSM265 (Figure 1C). While none of the mutations in DHODH conferred resistance to this compound, we previously reported that TCMDC-125334 appears to exclusively target the DHODH enzyme. Expression of the *S. cerevisiae* cytosolic type I DHODH (*Sc*DHODH) ablates the activity of TCMDC-125334, and treating the *Sc*DHODH line with TCMDC-125334 and proguanil in combination does not impact this activity^47,48^. In contrast, for compounds that target cytochrome bc1, proguanil has a potentiating effect in the *Sc*DHDOH line ^49^.

**Figure 1.**
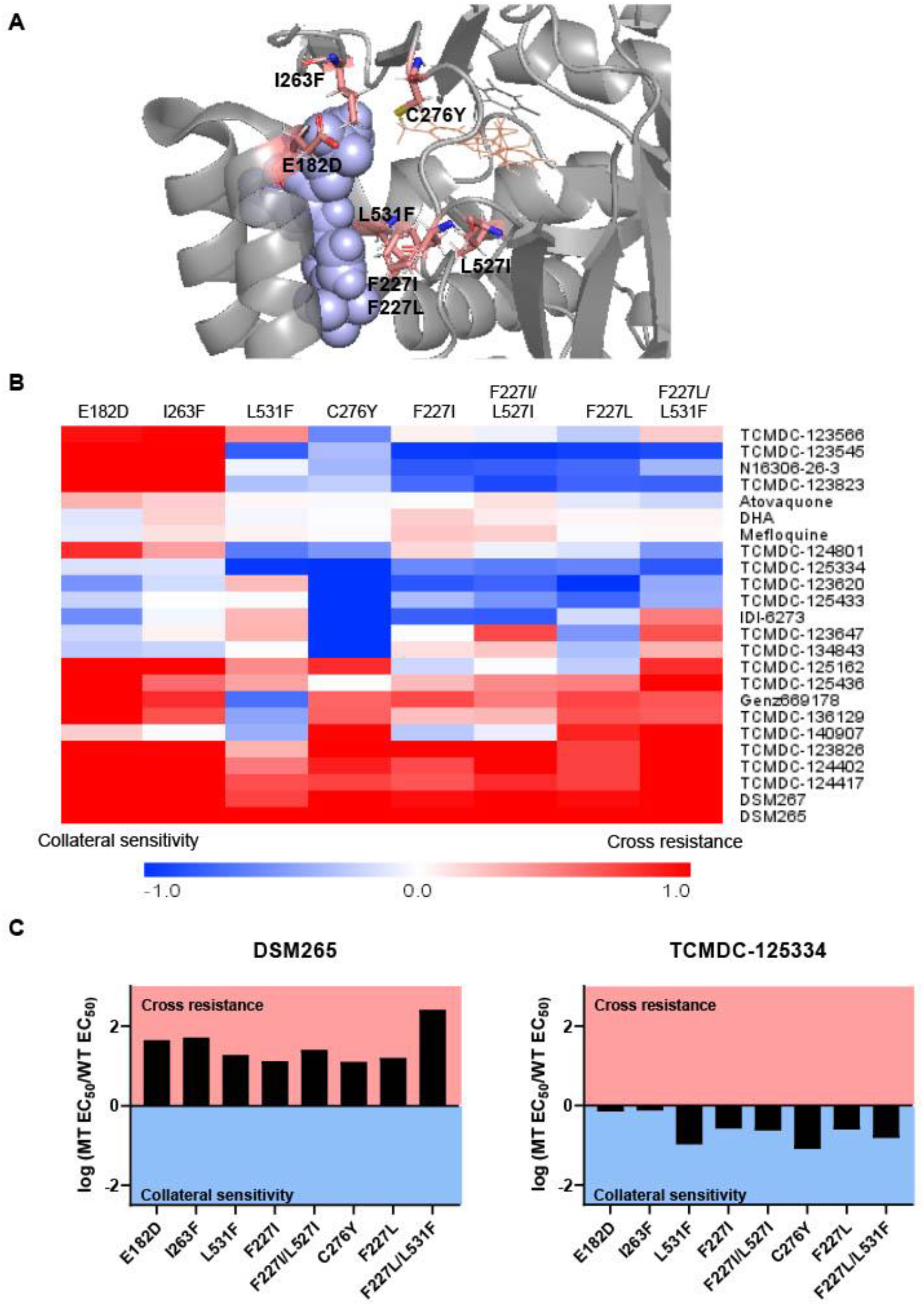
Identifying DHODH inhibitors with broad activity against mutant lines. We tested several DSM265-resistant parasite lines against a set of 17 compounds identified from a previous screen of GSK libraries^47^. A) Crystal structure of DHODH bound to DSM265 (PDB ID: 4RX0). Mutations included in our cross-resistance analysis are highlighted in pink B) To visualize patterns of cross-resistance and collateral sensitivity, we calculated the fold change of each mutant line over its wildtype parent, and plotted the log_10_-transformed values in a heatmap. Hierarchical clustering by Euclidean Distance based on both parasite line and compound was performed using MultiExperimentViewer v4.9. Shades of red indicate that a parasite line is resistant to the indicated compound while blue indicates that it is sensitive. Note that data from the DHODH E182D, L531F, I263F, F227I, and F227I/L531F mutant lines was previously reported^47^. C) One compound, TCMDC-125334 stands out in this analysis as being active against all mutant lines tested. Shown is a bar graph of the log fold change in EC_50_ relative to wildtype for all eight mutant lines tested against DSM265 and TCMDC-125334. DSM265 data was previously reported.^9^

### Resistance to TCMDC-125334 can be conferred by copy number variation as well as the novel point mutation DHODH I263S

Since TCMDC-125334 showed activity against all DHODH mutant lines tested, we wanted to characterize the evolution of resistance to this compound to determine its independent ability to select resistant mutants. We performed *in vitro* selections with three independent populations of 10^8^ *Pf*3D7 A10 parasites ^50^. Cultures were treated with 830nM TCMDC-125334 (approximately the EC_99_) until no living parasites were visible by thin smear microscopy, then allowed to recover in compound-free media. After two rounds of treatment, populations displayed moderate (∼2-fold) resistance to TCMDC-125334 (Figure 2A,B). Selected parasites were cloned by limiting dilution. Clonal lines isolated from flask 1 had two or three-fold copy number amplification encompassing the *dhodh* locus, which corresponded to a two or three-fold resistance phenotype to TCMDC-125334, as well as to the structurally-distinct DHODH inhibitors DSM265, Genz669178, and IDI6273. Of the clones isolated from flask 2, four did not have a resistance phenotype, and had no genetic changes in the *dhodh* locus (Figure 2C-F, Table 1, Figure S2, Data File S1, Data File S2, Data File S3, Table S2). One clone, T-F2-C1, was 3-fold resistant to TCMDC-125334 and Genz669178, and 13-fold resistant to IDI-6273 (Figure 2C-F, Table 1, Data File S1). Whole-genome sequencing (WGS) revealed that clone T-F2-C1 had a point mutation resulting in a DHODH I263S amino acid change (Data File S3, Table S2). Notably, this DHODH I263S parasite line remained sensitive to DSM265 (Figure 2D, Table 1, Data File S1).

**Figure 2.**
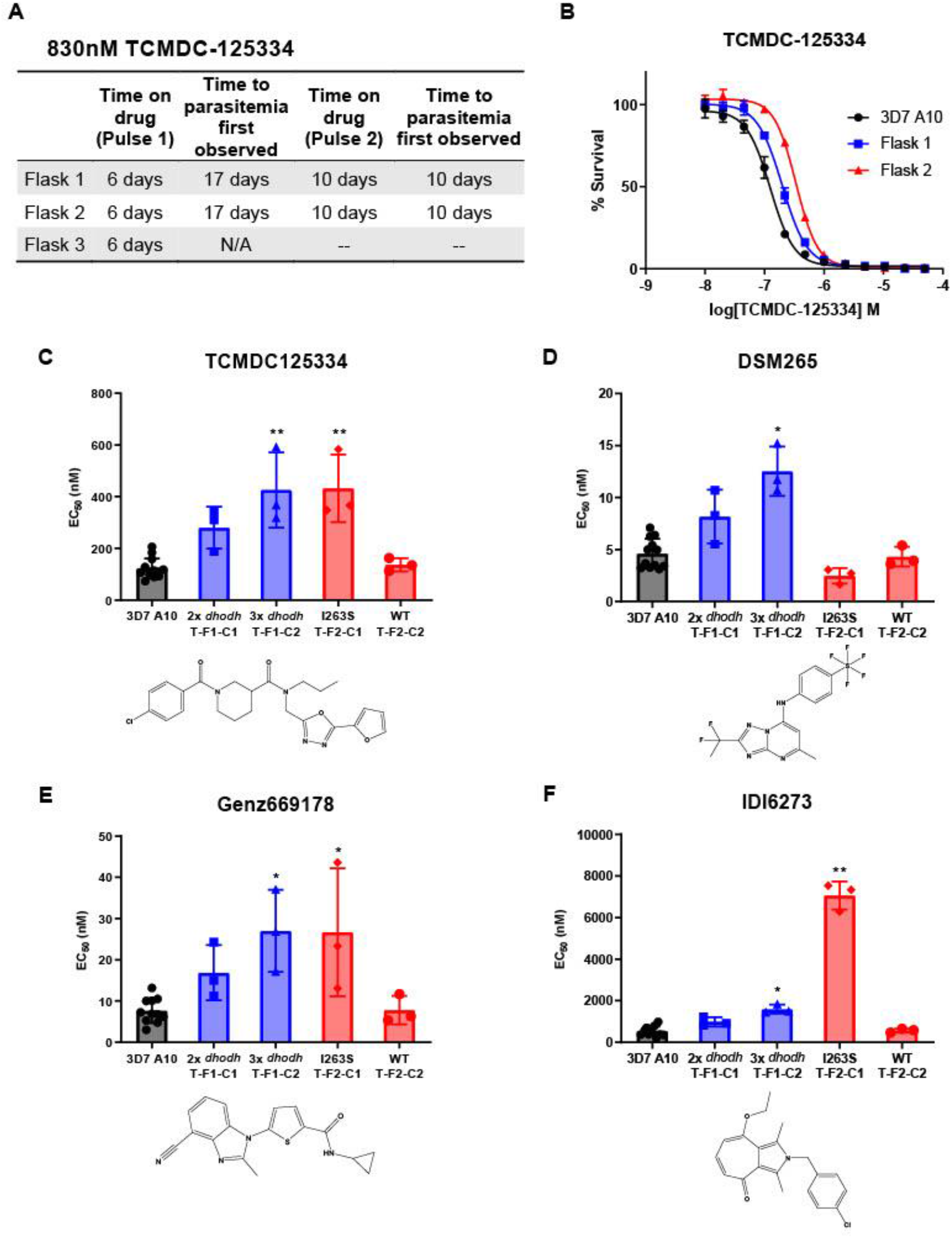
in vitro resistance to TCMDC125334 can be mediated by copy number variation as well as the novel point mutation DHODH I263S. A) Protocol for *in vitro* selection with TCMDC-125334. Parasite populations in three independent 25mL culture flasks were exposed to 830nM TCMDC-125334, then allowed to recover in the absence of drug. Resistant parasites emerged after two rounds of treatment within the indicated timeframe. B) *in vitro* dose response curve of bulk populations recovered after the second pulse of drug treatment. Error bars show standard deviation of technical replicates. C-E) Resistant populations were cloned by limiting dilutions. Shown is the average dose response phenotype of two representative clones from each flask for TCMDC-125334 (C), DSM265 (D), Genz669178 (E), and IDI-6273 (F). Bar graphs represent average EC_50_ with error bars depicting standard deviation. Individual bioreplicates are also shown. Compound structures for TCMDC-125334, DSM265, Genz669178, and IDI-6273 are displayed below their corresponding graphs. Statistical significance was determined by a Kruskal-Wallis test, with post-hoc multiple comparisons (Dunn’s) of each clone to 3D7 A10. *P ≤ 0.05; **P ≤ 0.01

**Table 1:** DHODH genotype and corresponding dose response phenotype for *in vitro* TCMDC-125334 selected lines.

| Clone ID | DHODH Mutation(s)<sup>1</sup> | DHODH CNV | TCMDC-125334 EC<sub>50</sub> (nM) | DSM265 EC<sub>50</sub> (nM) | IDI-6273 EC<sub>50</sub> (nM) | Genz669178 EC<sub>50</sub> (nM) |
| --- | --- | --- | --- | --- | --- | --- |
| 3D7 A10 | WT | 1 | 125±38.4 | 4.30±1.43 | 518±198 | 7.28±2.64 |
| T-F1-C1 | WT | 2 | 280±81.3 | 8.17±2.59 | 993±220 | 16.9±6.70 |
| T-F1-C2 | WT | 3 | 426±146** | 12.5±2.36* | 1597±212*10 | 27.0±10.0* |
| T-F1-C3 | WT | 2 | 298±36.5* | 8.71±0.370 | 1330±234 | 17.6±7.90 |
| T-F2-C1 | I263S | 1 | 432±130.8** | 2.47±0.740 | 7057±671** | 26.7±15.5* |
| T-F2-C2 | WT | 1 | 137±25.1 | 4.31±0.944 | 572±91.5 | 7.79±3.42 |
| T-F2-C3 | WT | 1 | 202±60.9 | 4.11±1.70 | 515±103 | 7.91±3.24 |
| T-F2-C4 | WT | 1 | 161±28.6 | 5.07±1.24 | 692±211.1 | 8.03±0.729 |
| T-F2-C5 | WT | 1 | 157±33.9 | 3.04±0.697 | 404±51.3 | 6.10±3.00 |
| Overall P-value (approximate) |  |  | 0.0004 | 0.0013 | 0.0007 | 0.0013 |
| Kruskal-Wallis Statistic |  |  | 28.52 | 25.54 | 27.05 | 25.50 |

Shown is mean EC<sub>50</sub> +/- standard deviation. Statistical significance was determined by a Kruskal-Wallis test, with post-hoc multiple comparisons (Dunn's) of each clone to 3D7 A10. \*P ≤ 0.05; \*\*P ≤ 0.01.
Overall statistics are reported for each comparison group.
<sup>1</sup>Variants identified by whole-genome sequencing

In a second independent biological replicate, we treated three additional parasite populations with 1µM TCMDC-125334. Similarly, bulk populations recovered after either one or two rounds of treatment displayed moderate (∼2-fold) resistance to multiple DHODH inhibitors, and exhibited copy number variation (CNV) at the *dhodh* locus as detected by qPCR (Figure S2, Figure S3). Two of the populations treated with 1µM were subjected to a second round of treatment at 1µM TCMDC-125334 for 13 days, then 1.5µM TMCDC-125334 for 15 days. These populations developed a stronger resistance phenotype (Figure S3), and correspondingly showed 8-fold increased copy number at the *dhodh* locus (Figure S2), while no mutations were detected in the *dhodh* locus by whole-genome sequencing (Data File S4, Table S2).

As a control, we performed a parallel set of selections with 30nM DSM265 (the EC_99_). As previously observed, parasites resistant to DSM265 emerged rapidly after just one round of treatment. WGS of the bulk populations revealed two point mutations in DHODH—DHODH C276Y and I273M (Figure S4, Table S2, Table S3, Data File S4).

### Sequential treatment of DSM265-resistant parasites with TCMDC125334 selects for additional mutations in DHODH

Given the hypersensitivity of DHODH C276Y parasites to TCMDC-125334, we also wanted to test what would happen if we treated DHODH C276Y parasites with this compound. In our previous work, selections with two distinct DHODH inhibitors, Genz666136 and DSM74, yielded resistant parasites with a DHODH E182D mutation, which were hypersensitive to the DHODH inhibitor IDI-6273. When we treated the DHODH E182D parasite line with IDI-6273, parasites reverted to the wildtype D182 protein sequence, albeit with a different DNA codon. The D182E revertant line had a drug sensitivity phenotype similar to wildtype parasites ^36^. We hypothesized that treating DHODH C276Y parasites with TCMDC-125334 would similarly cause a reversion to wildtype, and thus offer a strategy to regain sensitivity to DSM265 (Figure 3A).

**Figure 3.**
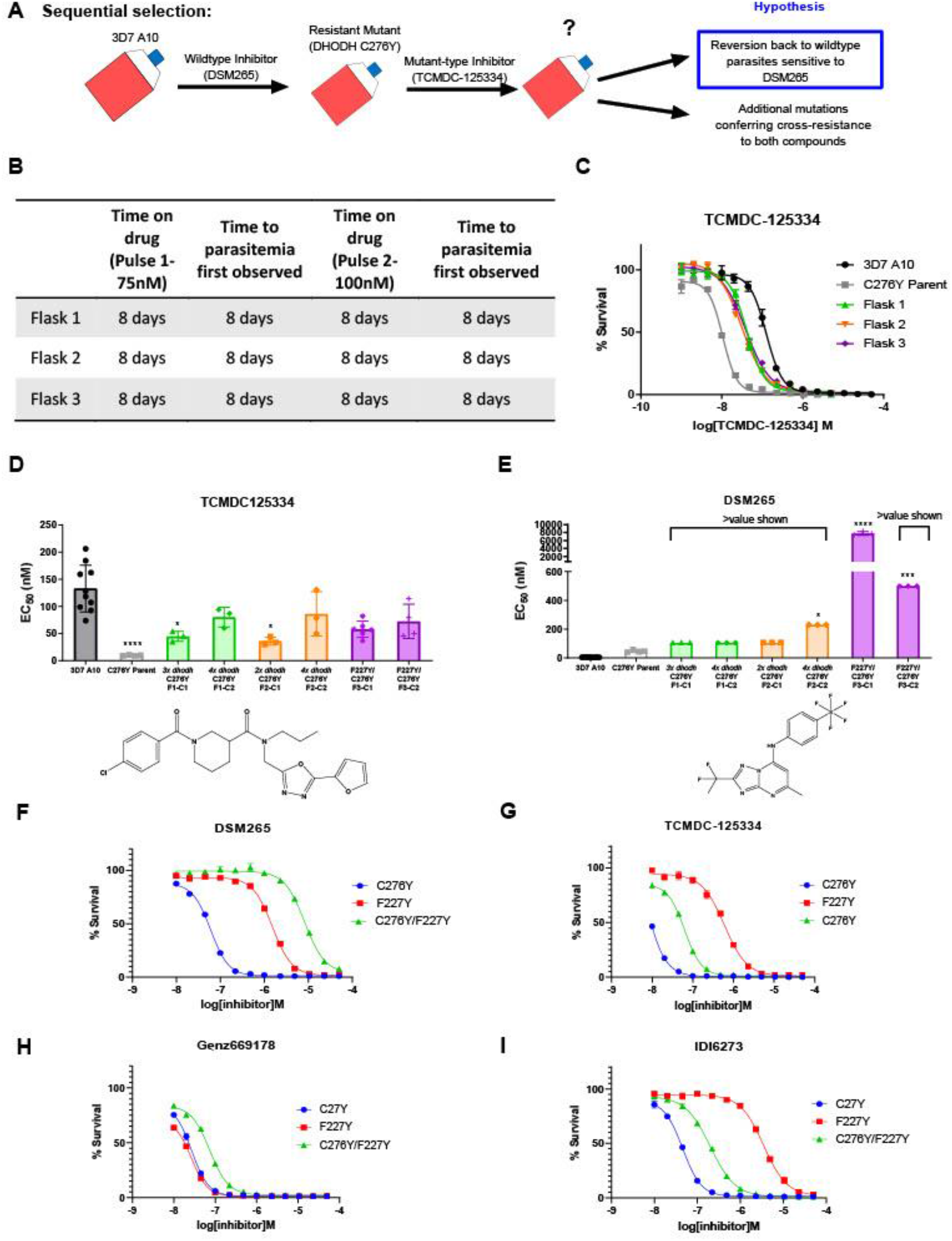
Mutant parasites can overcome collateral sensitivity by acquiring additional genetic changes that confer high-level resistance to wildtype inhibitors. A) General schematic for sequential selection. B) Protocol for *in vitro* selection. DHODH C276Y mutant parasites in three independent 25mL culture flasks were exposed to 40nM TCMDC-125334, then allowed to recover. The time on drug and time to recovery are indicated. After two rounds of treatment and recovery, resistant parasites are observed in all three flasks. C) *in vitro dose* response curve of selected bulk populations. Error bars show standard deviation of technical replicates. D-E) Resistant populations were cloned by limiting dilution. Shown is the dose response phenotype of two representative clones from each flask for TCMDC-125334 (C), and DSM265 (D), with chemical structures illustrated below. Bar graphs represent average EC_50_ with error bars depicting standard deviation. Individual bioreplicates are also shown. Statistical significance was determined by a Kruskal-Wallis test, with post-hoc multiple comparisons (Dunn’s) of each clone to 3D7 A10. In cases where the EC50 could not be determined with range of concentrations tested, the maximum concentration that achieved >50% growth was used as a stand-in value. *P ≤ 0.05; ***P≤ 0.001; ****P≤ 0.0001 F-I) The DHODH C276Y/F227Y double mutant exhibits a resistant phenotype intermediate to single mutations. The dose response phenotypes of th DHODH C276Y/F227Y line selected in this study (clone C276Y T-F3-C3), the DHODH C276Y line previously selected with DSM265, and the DHODH F227Y line previously selected with DSM265 (Mandt et al., 2019), were characterized. Shown is a representative dose response curve for DSM265 (F), TCMDC-125334 (G), Genz669178 (H), and IDI-6273 (I) for illustration. Error bars show standard deviation of technical replicates.

For these experiments, we used the DHODH C276Y clone that we had previously isolated from *in vitro* selections with DSM265 ^9^. We treated three 25mL flasks of approximately 10^8^ DHODH C276Y parasites with 75nM TCMDC-125334. After parasites recovered, populations were treated with a second round of 100nM TCMDC-125334. Resistance was observed in all three bulk parasite populations recovered from this second pulse (Figure 3B,C). While these parasites were less sensitive to TCMDC-125334 relative to the DHODH C276Y parental line, they were still more sensitive than *Pf*3D7 A10 parasites with a wildtype DHODH sequence (Figure 3C). Clones isolated from these sequentially selected populations had a similar intermediate dose response phenotype for TCMDC-125334 (Figure 3D, Table 2, Data File S1). However, compared to the DHODH C276Y parent, they had greatly increased resistance to DSM265, with EC_50_’s >200nM (Figure 3E, Table 2, Data File S1). Whole genome sequencing revealed that parasites retained the DHODH C276Y mutation, and gained additional mutations in DHODH, rather than reverting back to wildtype (Data File S2, Data File S3, Table S2). Clones isolated from flask 1 and flask 2 had increased *dhodh* copy number, in addition to the DHODH C276Y mutation, as confirmed by qPCR (Table 2, Figure S5, Data File S2). Clones from flask 3 had gained an additional F227Y mutation (Table 2, Data File S3, Table S2).

**Table 2:** DHODH genotype and corresponding dose response phenotype for *in vitro* selection of DHODH C276Y parent with TCMDC-125334.

| Clone ID | DHODH Mutation(s)<sup>1</sup> | DHODH CNV | TCMDC-125334 EC<sub>50</sub> (nM) | DSM265 EC<sub>50</sub> (nM) | IDI-6273 EC<sub>50</sub> (nM) | Genz669178 EC<sub>50</sub> (nM) |
| --- | --- | --- | --- | --- | --- | --- |
| 3D7 A10 | WT | 1 | 125±38.4 | 4.30±1.43 | 518±198 | 7.28±2.64 |
| C276Y Parent | C276Y | 1 | 9.44±0.798**** | 47.6±8.69 | 41.7±12.7**** | 22.0±6.68 |
| C276Y-T-F1-C1 | C276Y | 3-4 | 44.7±9.35* | >106 | 185±36.1 | >106* |
| C276Y-T-F1-C3 | C276Y | 4 | 45.9±11.2 | >158 | 199±58.6 | >129** |
| C276Y-T-F1-C2 | C276Y | 4 | 80.2±17.9 | >106 | 345±64.1 | >106** |
| C276Y-T-F1-C4 | C276Y | 2-3 | 54.5±21.0 | >106 | 191±95.4 | >230** |
| C276Y-T-F2-C1 | C276Y | 2 | 35.8±7.20** | >106 | 138±44.0* | >106 |
| C276Y-T-F2-C3 | C276Y | 2-3 | 43.0±7.81* | >106 | 164±12.0 | >106 |
| C276Y-T-F2-C4 | C276Y | 4 | 60.2±22.0 | >230** | 232.0±39.2 | >156**** |
| C276Y-T-F2-C2 | C276Y | 4 | 86.0±40.8 | >230* | 326±155 | >230*** |
| C276Y-T-F3-C1 | C276Y/F227Y | 1 | 57.7±14.8 | 7800±458**** | 163±36.3** | 69.7±14.5 |
| C276Y-T-F3-C2 | C276Y/F227Y | 1 | 81.1±32.0 | >499*** | 210±80.5 | 113±52.5 |
| C276Y-T-F3-C3 | C276Y/F227Y | 1 | 53.6±10.0 | >499*** | 174±23.74 | 90.5±34.4 |
| Overall P-value (approximate) |  |  | <0.0001 | <0.0001 | <0.0001 | <0.0001 |
| Kruskall-Wallis Statistic |  |  | 38.12 | 47.36 | 45.55 | 54.32 |

Shown is mean EC<sub>50</sub> +/- standard deviation. Statistical significance was determined by a Kruskal-Wallis test, with post-hoc multiple comparisons (Dunn's) of each clone to 3D7 A10. In cases where the EC<sub>50</sub> could not be determined with range of concentrations tested, the maximum concentration that achieved >50% growth was used as a stand-in value. \*P ≤ 0.05; \*\*P ≤ 0.01; \*\*\*P ≤ 0.001; \*\*\*\*P ≤ 0.0001. Overall statistics are reported for each comparison group.
<sup>1</sup>Variants identified by whole-genome sequencing

We had previously selected the DHODH F227Y single mutation with the triazolopyrimidine-based inhibitor DSM267, although we had not tested this mutant line against the compounds in the Tres Cantos Antimalarial Set. We characterized the dose-response phenotype of the DHODH F227Y mutant line and found that it exhibited 3-fold resistance to TCMDC-125334 (Table 3, Data File S1). Comparing the dose response phenotype of the DHODH C276Y/F227Y double mutant with the dose response of the two single mutant lines DHODH C276Y and DHODH F227Y, we find that the phenotype of the two mutations in combination is consistent with additivity, ruling out a negative epistatic interaction (Figure 3 F-I, Table 3, Data File S1). Overall, sequential selection of a DSM265-resistant line with TCMDC-125334 does not restore sensitivity to DSM265. While sequentially-selected parasites with additional mutations in DHODH were highly-resistant to DSM265, they were not cross-resistant to TCMDC-125334, suggesting that they would still be sensitive to a therapeutic dose.

**Table 3:** Resistance phenotype of C276Y and F227Y single mutants and C276Y/F227Y double mutant compared to expected phenotype under additive epistasis.

| **DHODH Mutations** | **TCMDC-125334**<br>**EC<sub>50</sub> (fold**<br>**change)** | **DSM265 EC<sub>50</sub>**<br>**(fold change)** | **IDI-6273 EC<sub>50</sub>**<br>**(fold change)** | **Genz669178**<br>**EC<sub>50</sub> (fold change)** |
| --- | --- | --- | --- | --- |
| **3D7 A10** | 125±38.4 | 4.30±1.43 | 518±198 | 7.28±2.64 |
| **C276Y** | 9.44±0.798 (0.07) | 47.6±8.69 (11.1) | 41.7±12.7 (0.081) | 22.0±6.68 (3.02) |
| **F227Y** | 408±17.6 (3.3) | 998±50.0 (232) | 2210±289 (4.3) | 18.8±3.39 (2.6) |
| **C276Y/F227Y (F3B6)** | 57.7±14.8 (0.46) | 7801.0±457.9<br>(1814) | 163.3±36.3 (0.32) | 69.7±14.5 (9.6) |
| **Expected fold change**<br>**for C276Y/F227Y**<br>**(additive assumption)** | 0.23 | 2575 | 0.35 | 7.9 |

EC<sub>50</sub> (nM) of each line is reported, with fold change relative to 3D7 A10 in parentheses. The expected fold change of the C276Y double mutant based on an assumption of additive epistasis is reported for comparison.

### Treatment with TCMDC125334 + DSM265 combination delays but does not suppress resistance

Based on the collateral sensitivity observed between DSM265 and TCMDC-125334, we hypothesized that treatment with a combination of DSM265 and TCMDC-125334 would suppress the emergence of resistant parasites (Figure 4A). We treated three independent populations of 10^8^ *Pf*3D7 A10 parasites with both compounds simultaneously, using the EC_99_ of each. After three pulses of drug treatment totaling 63-73 days, we obtained two populations cross-resistant to both compounds (Figure 4B,C). Three of the four clonal lines isolated from Flask 1 were moderately (∼2-fold) resistant to both DSM265 and TCMDC-125334, as well as Genz669178 and IDI-6273 (Figure 4D-G, Table 4, Data File S1). Copy number duplication of the *dhodh* locus was detected in these resistant clones by qPCR (Table 4, Figure S6). All six clones isolated from Flask 2 were 4-to-6-fold resistant to TCMDC-125334 and 16-20-fold resistant to DSM265. These clones were also ∼2-fold resistant to IDI-6273, but were still sensitive to Genz669178 (Figure 4D-G, Table 4, Data File S1). Sequencing revealed that these six clones had the same point mutation resulting in a DHODH V532A amino acid change (Data File S3, Data File S5, Table S2). Thus, although resistance took longer to emerge with combination treatment compared to treatment with DSM265 or TCMC-125334 alone, this strategy was ultimately insufficient to suppress the emergence of parasites resistant to both compounds. This result would argue against using these compounds in combination.

**Figure 4.**
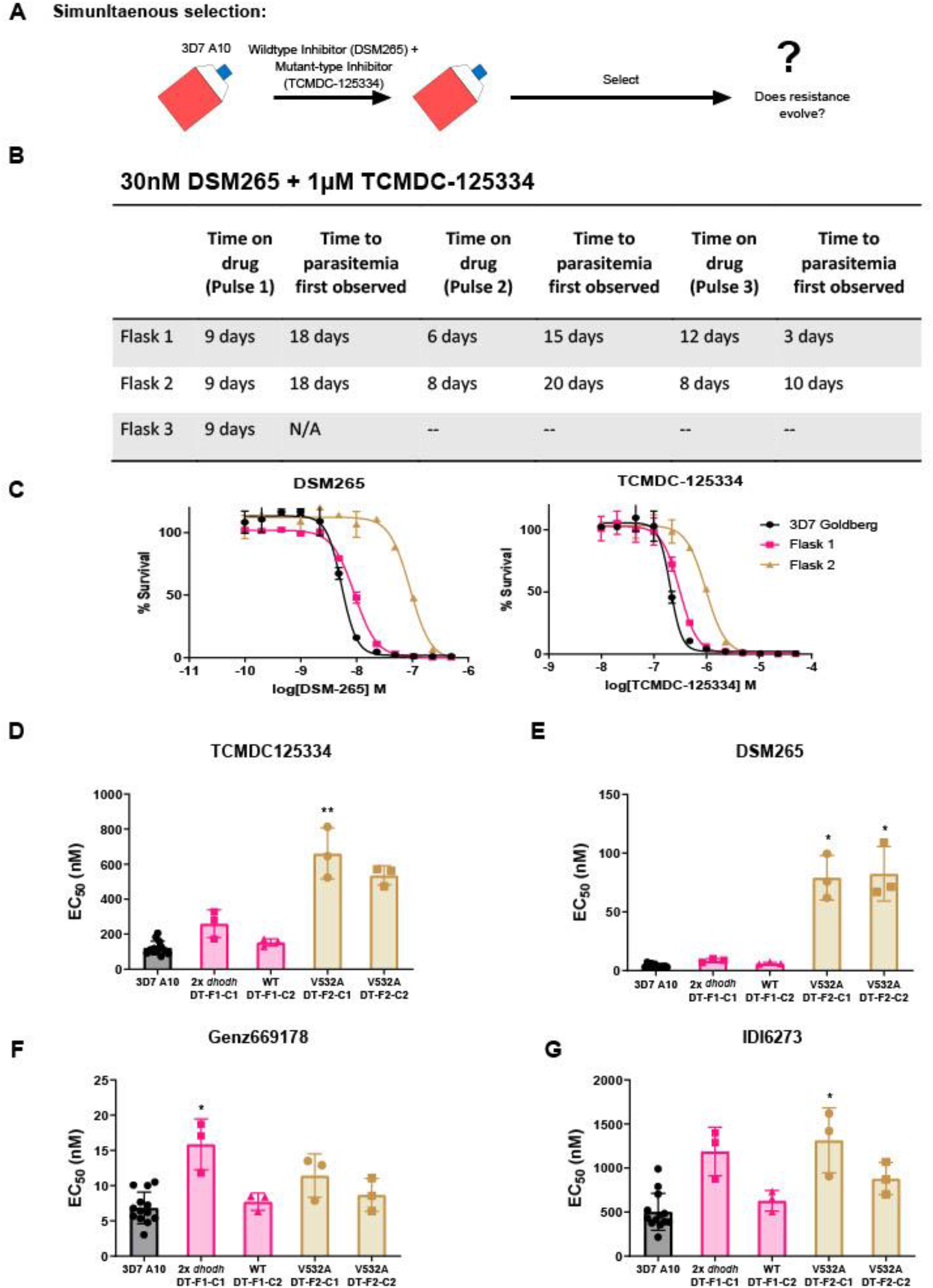
Resistance to the DSM265+TCMDC-125334 combination arose after three rounds of treatment, with cross-resistance to both compounds conferred by the novel DHODH V532A mutation. A) General schematic for simultaneous selection. B) Protocol for *in vitro* selection with TCMDC-125334 and DSM265. Parasite populations in three independent 25mL culture flasks were exposed to combination treatment, then allowed to recover. The time on drug and time to recovery are indicated. After three rounds of treatment and recovery, resistant parasites are observed in Flasks 1 and 2. C) *in vitro* dose response curve of bulk populations from flasks 1 and 2. Error bars show standard deviation of technical replicates. D-G) Resistant populations were cloned by limiting dilutions. Shown is the dose response phenotype of two representative clones from each flask for TCMDC-125334 (D), DSM265 (E), Genz669178 (F), and IDI6273 (G) with chemical structures of each compound illustrated below. Bar graphs represent average EC_50_ with error bars depicting standard deviation. Individual bioreplicates are also shown. Statistical significance was determined by a Kruskal-Wallis test, with post-hoc multiple comparisons (Dunn’s) of each clone to 3D7 A10. *P ≤ 0.05; **P ≤ 0.01

**Table 4:** DHODH genotype and corresponding dose response phenotype for in vitro TCMDC-125334 + DSM265 selected lines.

| Clone ID | DHODH Mutation(s) | DHODH CNV | TCMDC-125334 EC<sub>50</sub> (nM) | DSM265 EC<sub>50</sub> (nM) | IDI-6273 EC<sub>50</sub> (nM) | Genz669178 EC<sub>50</sub> (nM) |
| --- | --- | --- | --- | --- | --- | --- |
| 3D7 A10 | WT | 1 | 125±38.4 | 4.30±1.43 | 518±198 | 7.28±2.64 |
| DT-F1-C1 | WT<sup>1</sup> | 2 | 260±79.5 | 8.59±1.60 | 1189±276 | 15.9±3.61* |
| DT-F1-C2 | WT<sup>2</sup> | 1 | 153±21.0 | 6.09±0.875 | 628±117 | 7.74±1.23 |
| DT-F1-C3 | WT<sup>1</sup> | 2 | 191±64.0 | 7.51±0.076 | 895±349 | 11.0±3.74 |
| DT-F1-C4 | WT<sup>2</sup> | 2 | 319±137 | 11.2±4.50 | 1649±655* | 19.5±8.39* |
| DT-F2-C1 | V532A<sup>1</sup> | 1 | 661±145** | 79.2±19.0* | 1316±368* | 11.4±3.08 |
| DT-F2-C2 | V532A<sup>1</sup> | 1 | 536±54.4 | 82.4±23.1* | 881±185 | 8.69±2.35 |
| DT-F2-C3 | V532A<sup>2</sup> | 1 | 816±480* | 93.2±36.9** | 1377±669* | 12.8±6.32 |
| DT-F2-C4 | V532A<sup>2</sup> | 1 | 676±164** | 94.2±17.2** | 1328±410* | 12.0±3.67 |
| DT-F2-C5 | V532A<sup>2</sup> | 1 | 556±142* | 73.4±16.4* | 1029±347 | 9.13±1.45 |
| DT-F2-C6 | V532A<sup>2</sup> | 1 | 674±118** | 87.6±25.3** | 1307±412* | 14.4±4.16 |
| Overall P-value (approximate) |  |  | <0.0001 | <0.0001 | 0.0019 | 0.0091 |
| Kruskall-Wallis Statistic |  |  | 35.72 | 35.80 | 27.85 | 23.50 |

Shown is mean EC<sub>50</sub> +/- standard deviation. Statistical significance was determined by a Kruskal-Wallis test, with post-hoc multiple comparisons (Dunn's) of each clone to 3D7 A10. \*P ≤ 0.05; \*\*P ≤ 0.01.
Overall statistics are reported for each comparison group.
WT= Wildtype
<sup>1</sup>DHODH genotype determined by whole-genome sequencing
<sup>2</sup>DHODH genotype determined by Sanger sequencing

### *in silico* docking of TCMDC-125334 to *Pf*DHODH suggests molecular mechanisms of resistance

To better understand the binding mode of TCMDC-125334 and develop inferences on the molecular mechanisms by which mutations characterized in this study may confer resistance or hypersensitivity to the compound, we performed *in silico* molecular docking using Flare™, v.4.0.3.40719 (Cresset-Group). The co-crystal structures of DHODH, FMN, orotate, and a variety of inhibitors have previously been reported, but there is no reported structure with TCMDC-125334^10,11,46,51^. The best pose predicted by our simulations shows that TCMDC-125334 docks within same hydrophobic binding pocket as DSM265 and other DHODH inhibitors (Figure 5A&B). As with all published crystal structures of DHODH inhibitors, our simulations suggest that TCMDC-125334 binds allosterically with regards to both orotate/dihydroorotate and FMN.

**Figure 5.**
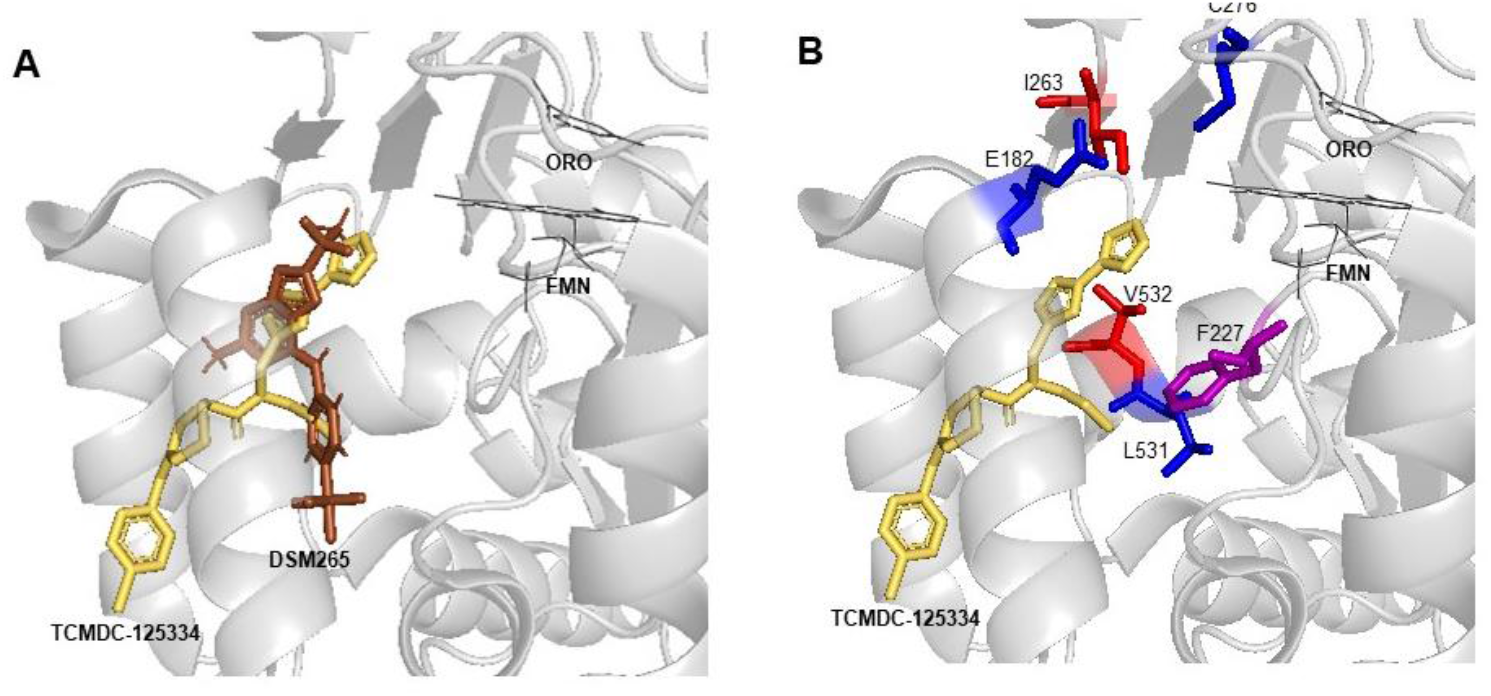
Molecular docking of TCMDC-125334 with *Pf*DHODH. A) TCMDC-125334 docks within the inhibitor binding pocket of DHODH. Shown is overlay of TCMDC-125334 docked in combined structure, and DSM265 from the 4RX0 structure. B) Mutations conferring cross-resistance or collateral sensitivity are in close proximity to TCMDC-125334. Key residues discussed in the study are highlighted. Residues conferring cross-resistance are shown in red, while residues conferring collateral sensitivity are in blue. The F227 residue is colored purple, as the F227L and F227I substitutions confer collateral sensitivity while the F227Y substitution confers resistance.

We next wanted to use our modeling results to explore how mutations in DHODH might impact resistance at a molecular level. Our docking results suggest that the DHODH^V532A^ mutation would disrupt hydrophobic interactions with the furan and 1,3,4-oxadiazole rings, consistent with the observed 5.3-fold resistance seen with the DHODH^V532A^ mutants. There are no published crystal structures for the DHODH^C276Y^ so we used Flare’s mutate function to produce a model of DHODH^C276Y^, optimized the results in the absence of other molecules and then in the presence of TCMDC-125334, orotate, and FMN with Flare’s protein minimize function, and rescored the result. Our simulation predicted that the DHODH^C276Y^ mutant would have higher affinity than wildtype.

### Cross-resistant parasites are relatively fit in *in vitro* growth assays

Given that the DHODH V532A mutation confers cross-resistance to DSM265 and TCMDC-125334, we wanted to assess the relative fitness of *P. falciparum* parasites carrying this mutation. Previously we have shown that some DHODH mutations, such as DHODH E182D, can negatively impact enzyme function and *in vitro* growth, while other mutations, such as DHODH C276Y do not appear to confer a fitness cost ^9,46^. Because resistance to simultaneous selection with DSM265 and TCMDC125334 took longer to emerge, we hypothesized that DHODH V532A mutants might have a growth defect relative to wildtype parasites. To test this hypothesis, we performed *in vitro* competitive growth assays. Synchronized DHODH V532A and wildtype 3D7 A10 lines were mixed in a 1:1 ratio. The mixed culture was grown in three independent flasks over a four-week period in drug-free media, and genomic DNA was collected every 7-9 days for one month. The ratio of mutant and wildtype parasites was determined by the calculating the percentage of reads detecting the V532A allele in whole-genome sequencing. This analysis showed that the DHODH V532A variant remained at ∼50% frequency over time (Figure 6, Table S4). As a confirmation of this result, we also assessed the resistance phenotype of mixed cultures. On Day 0, the mixed culture exhibited an intermediate dose response phenotype, as illustrated by a biphasic dose-response curve. This intermediate phenotype was maintained at Day 28 (Figure S7). Overall, both the genotypic and phenotypic data indicate that the DHODH V532A mutant is as fit as wildtype parasites.

**Figure 6.**
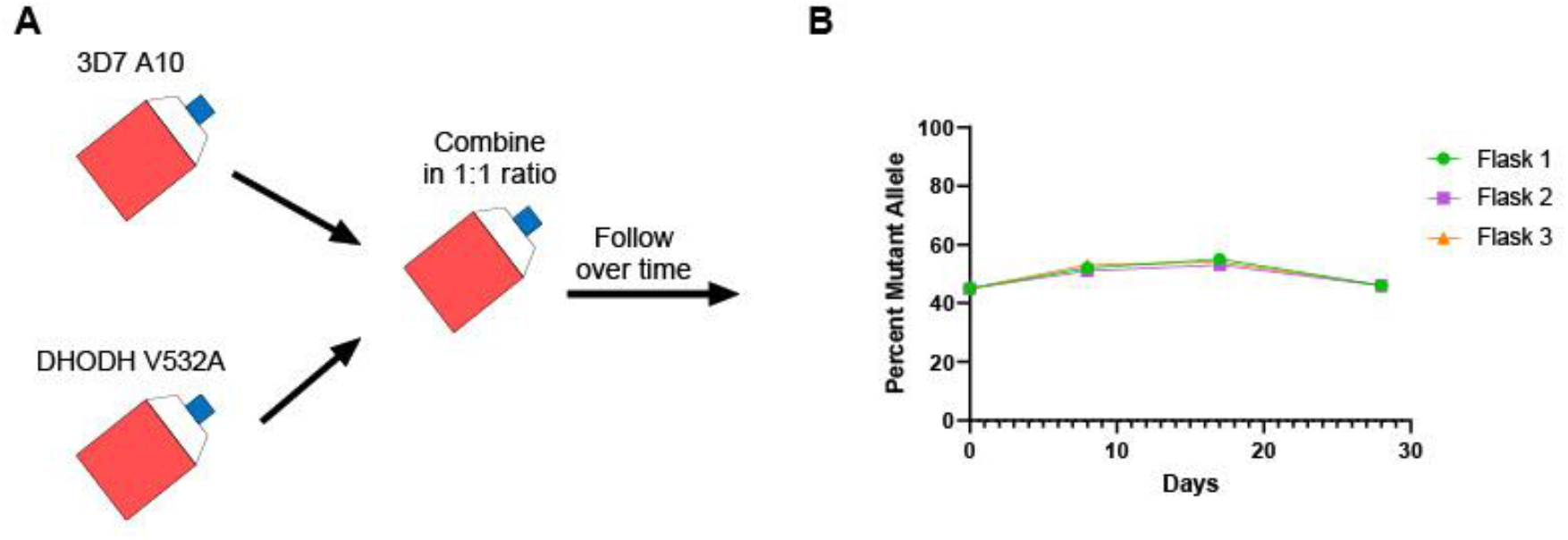
The DHODH V532A mutation does not confer a fitness cost in in vitro competitive growth assays. A) Schematic of competitive growth experiments. Synchronized 3D7 parent and DHODH V532A mutant parasites at 1% ring stage parasitemia were mixed at equal ratios. The mixed culture was then split into three independent flasks, and followed over time. Mixed cultures were grown for four weeks, and genomic DNA was collected every 7-9 days. B) The percent mutant allele was calculated at each timepoint based on whole-genome sequencing reads.

## Discussion

The main conclusion of our study is that combining two DHODH inhibitors based on their collateral sensitivity profiles fails to suppress the emergence of resistance in *Plasmodium falciparum in vitro*. Although all previously-identified mutations that we tested exhibited sensitivity to TCMDC-125334, we identified the point mutation DHODH I263S in selection with TCMDC-125334 alone, which was resistant to TCMDC-125334 but not DSM265. We found that sequentially treating a collaterally sensitive mutant line with TCMDC-125334 selected for parasites with additional genetic changes in DHODH, including the DHODH F227Y mutation, which alone also confers resistance to TCMDC-125334. While the DHODH C276Y/F227Y double mutant is less sensitive than the DHODH C276Y parent, it is still more sensitive to TCMDC-125334 compared to wildtype parasites. Finally, we found that treatment with DSM265 and TCMDC-125334 in combination selected for cross-resistant parasites with a DHODH V532A mutation. We also demonstrated that the DHODH V532A mutation does not exhibit a fitness cost in *in vitro* competitive growth assays.

While we identified three point mutations in this study that confer resistance to TCMDC-125334, the majority of DHODH mutations tested thus far are sensitive to this molecule. Copy number variation (CNV) was a common mechanism of resistance to TCMDC-125334 across multiple independent selections. Three of four populations of wildtype 3D7 A10 selected with TCMDC-125334 exhibited increased copy number of DHODH. When we selected the collaterally sensitive DHODH C276Y line with TCMDC-125334, clones isolated from two of three independent populations exhibited 2 to-4-fold copy number variation of *dhodh*. In contrast to this, the majority of *in vitro* selections with DSM265 and DSM267 selected for point mutations in the *dhodh* locus, with copy number variation occurring only seldomly ^9^. While copy number variation confers cross-resistance across various DHODH inhibitor compound classes, this mechanism confers only moderately reduced susceptibility. CNVs are also less evolutionarily stable compared to point mutations. Others have shown that additional copies of *dhodh* are lost in *in vitro* culture over time ^52^. CNVs can also be lost during meiotic recombination in the mosquito stages.

Our finding that sequentially treating the DHODH C276Y line with a second inhibitor led to the acquisition of additional genetic changes, rather than reversion back to wildtype, has important implications for treatment strategies to manage resistance. This result is in contrast with our previous work, in which selecting DHODH E182D parasites with the mutant-type inhibitor IDI-6273 caused parasites to revert back to the wildtype amino acid sequence. Interestingly, we also observed that the DHODH E182D parasite had a competitive fitness defect ^36,46^, while we previously showed that the DHODH C276Y is as fit as wildtype in *in vitro* competitive growth assays ^9^. One caveat is that we have only looked at fitness in the blood stages both *in vitro* and *in vivo*. It is possible that mutations would affect the fitness of the parasites in mosquito or liver stages. Our results suggest that the fitness costs of resistance influence the evolutionary outcome of sequential treatment with a mutant-type inhibitor. In this case, a competitively fit mutant line favored alternative pathways to cross-resistance rather than reversion to wildtype. Similar research on collateral sensitivity in *Pseudomonas aeruginosa* from Barbosa et al. suggests that differing evolutionary outcomes (reversion vs. secondary mutation) may be due to differences in the fitness cost of the initial resistance mutations ^53^. Our work has implications for strategies such as drug cycling or the simultaneous use of multiple first line treatments that aim to delay the emergence of resistance ^54,55^. Such strategies assume that resistance mutations will disappear in the absence of pressure from the original selecting agent. However, our results suggest that if a resistant mutant is competitively fit, treatment with a second drug will more likely result in the acquisition of additional genetic changes and subsequent multidrug resistance.

Comparing the phenotype of the DHDOH C276Y/F227Y double mutant with DHODH C276Y and DHODH F227Y parasite lines, we also note that the two mutations in combination exhibit additive epistasis. This resulted in markedly high-level resistance to DSM265 (>1000-fold relative to wildtype). Multiple studies indicate that epistasis is an important factor in determining the evolution of resistance and cross-resistance/collateral sensitivity ^26,53,56-58^. Negative epistasis between resistance mutations within the same enzyme constrains the number of pathways to higher-level resistance ^57,58^. In the context of collateral sensitivity, the study from Barbosa et al. also indicates that re-sensitization to wildtype is favored when there is negative epistasis between resistance mechanisms ^53^. In contrast, our result highlights how, under additive epistasis, the emergence of an initial resistance mutation can open up pathways to higher-level resistance.

The major take-away of this study is that combination treatment with DSM265 and TCMDC-125334 failed to suppress resistance, and we hypothesize that this is due to the great diversity of evolutionarily competitive DHODH mutations that confer resistance to various inhibitors. Other reports in bacteria have also found that observed patterns of collateral sensitivity are often thwarted due to the existence of diverse trajectories to resistance. When selections are repeated enough times ^59^, or are performed with a large enough starting populations ^60^, cross-resistant variants eventually emerge, even if they are relatively rare. Similarly, we find that although most of the mutations we originally identified were collaterally-sensitive to TCMDC-125334, repeated treatment with DSM265 and TCMDC-125334 in combination ultimately yielded the cross-resistant variant DHDOH V532A.

This study also highlights the flexibility of the DHODH enzyme. Prior to this work, we had identified 13 individual point mutations and two sets of double mutations that conferred resistance to DHODH. In this study, we selected two additional point mutations—DHODH I263S and DHODH V532A, as well as the DHODH C276Y/F227Y double mutant line. Interestingly, the DHODH I263S mutation is not resistant to DSM265, while the DHODH I263F mutation at the same site confers strong (∼100-fold) resistance to this compound. Previous studies have crystallized the DHODH enzyme bound to various inhibitors, including DMS265 (PDB 4RX0), Genz669178 (PDB 4CQ8), and IDI-6273 (PDB 4CQA) ^8,36^. These compounds all bind within the same flexible pocket (see Figure 1A) of the enzyme. Our results demonstrate that this pocket not only accommodates the binding of a variety of chemical structures, but is also very mutationally flexible, highlighting the resistance liability of this enzyme as a drug target. Molecular simulation studies provided insight into the potential mechanism of resistance to both DSM265 and TCMDC-125334, highlighting the interactions at position V532. However, understanding the mechanism of collateral sensitivity will require further study. Molecular simulation points to differential binding affinity of TCMDC-125334 to the wildtype and C276Y mutant proteins, consistent with the increased sensitivity of the mutant observed in our study.

The *in vitro* studies described here also provide a framework for future *in vivo* work. We previously demonstrated that with DSM265 alone, the *in vitro* data is a good predictor of the *in vivo* evolution of resistance^9^. It will be important to test potential combination strategies *in vivo*, as pharmacokinetics and pharmacodynamics may impact how multiple drugs interact with each other, with implications for the emergence of resistance. However, our continued identification of novel mutations in the *dhodh* locus suggests that we still have not saturated the DHODH resistome. One implication of these results is that targeting this enzyme with antimalarial drugs has a high risk of resistance emergence, and brings into question the usefulness of pursuing further DHODH inhibitors.

## Methods

### Reagents and parasite strains

Atovaquone, dihydroartemisinin and mefloquine were purchased from Sigma-Aldrich (St. Louis, MO). IDI-6273 was purchased from ChemDiv (San Diego, CA). Genz669178 was graciously provided by Genzyme, a Sanofi Company (Waltham, MA). DSM265 was a kind gift from Margaret Phillips of Universitsy of Texas Southwestern (Dallas, TX). The *Pf*3D7 A10 clone used in this study was adapted to a 40 hour replication cycle in *in vitro* culture, and has been previously used in large-scale drug selection and sequencing efforts ^50^. Parasites with point mutations in *Pf*DHODH derived from previous work were generated as described ^9,46^. Compounds from GSK libraries were previously published in Ross et al., 2018^47^.

### *In vitro* parasite culturing

Parasites were cultured by standard methods with RPMI 1640 (Life Technologies) in 5% O^+^ human blood obtained from Interstate Blood Bank. RPMI 1640 was supplemented with 28mM NAaHCO_3_, 25mM HEPES, 50mg/mL hypoxanthine, 2525μg/mL gentamycin, and 0.5% Albumax II (Life Technologies). Parasite cultures were maintained in a gas mixture of 95% N_2_, 4% CO_2_, and 1.1% O_2_ at 37°C. Parasites were regularly synchronized by 5% sorbitol treatment ^61^, and all lines generated were preserved in glycerol stocks stored in liquid nitrogen.

### Resistance selections

*In vitro* resistance selections were performed in triplicate 25mL flasks with 3-4% ring-stage parasitemia. Selected populations were treated with the indicated dose of compound in RPMII daily until parasites were no longer visible by thin smear microscopy. Cultures were then replenished with compound-free RPMI, and monitored every 2-5 days by microscopy. Bulk recrudescent cultured were phenotyped in dose response assays. Resistant populations were cloned by limiting dilution in 96-well plates at a density of 0.2 parasites per well in the absence of compound.

### Dose response assays

To assess drug susceptibility phenotypes of parasite lines, ring-stage parasites were grown at 1% hematocrit, 1% starting parasitemia in 40µL volumes in 384-well black clear-bottomed plates. Parasites were cultured in the presence of test compounds plated in triplicate concentrations in twelve-point serial dilutions, and parasite growth after 72 hours was measured by a SYBR Green-based fluorescence assay ^62,63^. Lysis buffer with SYBR Green I at 10x concentration was added to the plates and allowed to incubate at room temperature for at least 12 hours. Fluorescence was measured at 494nm excitation, 530nm emission. Data was analyzed with CCD Vault, which calculates EC_50_ values based on a non-linear dose response curve fit (Burlingame, CA).

### Genomic DNA analysis

For genetic characterization of resistant clones, red blood cells infected with late-stage parasites were washed with 0.1% saponin to obtain isolated parasite pellets. For competitive growth assays, ring-stage infected red blood cells were directly suspended in DNA/RNA Shield (Zymo Research). Genomic DNA was then extracted using a DNAeasy Blood and Tissue Kit (Qiagen). ***Whole genome sequencing***. Libraries were prepared with the standard dual index protocol of the Nextera XT kit (Cat. No FC-131-1024). Whole-genome sequencing was performed on an Illumina NovaSeq 6000 with S4 200 chemistry and 100bp paired end reads. The *Plasmodium falciparum* 3D7 genome (PlasmoDB v. 13.0) was used as a reference for read alignment. Variants were called using an established analysis pipeline as previously described, where mutations are identified that are present in drug-selected parasites that are not in the drug-sensitive parent^50^. ***PCR and sanger sequencing***. PCR amplification of the *dhodh* locus was performed using Phusion High-Fidelity PCR Master Mix (New England BioLabs) as per the protocol. The locus was amplified in three overlapping segments with primers listed in Table S5. ***Quantitative PCR to assess copy number variation***. Copy number of the *dhodh* locus was assessed by quantitative PCR as previously described^46^, with power SYBR Green Master Mix (Applied Biosystems) and 0.1 ng gDNA template. qPCR was performed on a ViiA7 real-time PCR system with 384-well block (Applied Biosystems), and relative copy number was calculated using the ΔΔC_T_ method.

### *In silico* molecular modeling

For the modeling simulations conducted with Flare v.4.0.3.40719 (Cresset-Group™), the DHODH crystal structures (PDB codes: 4RX0, 3O8A, 4CQ8, 4CQA, and 4CQ9) were imported from the PDB and “Protein Prep” was performed using the default settings. Sequences were aligned and all structures superimposed to 4RX0. Three forms of TCMDC-125334 (unspecified enantiomer, R-isomer, and S-isomer) were then docked to all sequences using ensemble docking with the docking grid defined as all residues. Docking was performed using “Very Accurate but Slow” settings modified to allow rotation about amide bonds. Mutant structures were prepared with Flare’s mutate function using default settings, energy minimized with Flare’s protein minimize function in the absence of other molecules and then in the presence of TCMDC-125334, orotate, and FMN, and then rescored as described above. Simulations were visualized using PyMOL v. 4.6.0 (Intel).

### Competition growth assays

Mutant and wildtype cultures were synchronized at least twice prior to the assay. Ring-stage mutant and wildtype parasites at 5% hematocrit and 1% parasitemia were combined at equal volumes. Mixed culture was split into three 10mL replicate flasks, which were maintained for four weeks. Genomic DNA samples were collected and dose response assays performed at indicated timepoints.

### Statistical analysis

The heatmap visualizing patterns of cross-resistance and collateral sensitivity was created using MultiExperimentViewer (MeV) version 4.9.0. Hierarchical clustering of both parasite lines and compounds was performed based on Euclidean Distance using average linkage. Prism v8 (GraphPad) was used to make graphs and to perform statistical analysis comparing EC_50_’s of selected parasite clones and wildtype 3D7 A10. Significance was determined using a non-parametric ANOVA test (Kruskall-Wallis) with post-hoc multiple comparisons (Dunn’s test).

## Supporting information

Supplemental Materials (All Supplemental Figures and Tables))

## Data Availability

The raw whole-genome sequencing data generated in this study have been submitted to the NCBI Sequence Read Archive database (https://www.ncbi.nlm.nih.gov/sra/) under accession number PRJNA689594. Sanger sequencing of the PCR amplified *dhodh* locus have been submitted to GenBank (NCBI) under accession numbers MZ571149-MZ571158.

## Acknowledgements

We thank M. Phillips (UT Southwestern) and J. Burrows (Medicines for Malaria Venture) for generously providing compounds DSM265 and DSM267. We are also grateful to P. Hinkson (Harvard T.H. Chan School of Public Health) for technical support. We are grateful for generous financial support by the NIH (grant no. R01 AI093716 to D.F.W.), the Bill and Melinda Gates Foundation (Grand Challenges Exploration grant no. OPP1132451 to D.F.W., A.K.L., and FJ.G.), and the Harvard Malaria Initiative with support from ExxonMobil Foundation (to D.F.W.). R.E.K.M. was additionally supported by the Harvard Herchel Smith Fellowship, an NIH T32 grant, and funds from the ExxonMobil Foundation. M.R.L. was supported in part by a Ruth L. Kirschstein Institutional National Research Service Award T32 GM008666 from the National Institute of General Medical Sciences. This publication includes data generated at the UC San Diego IGM Genomics Center utilizing an Illumina NovaSeq 6000 that was purchased with funding from a National Institutes of Health SIG grant (#S10 OD026929).

## Author contributions

R.E.K.M. performed the dose response assays, *in vitro* selection experiments, and competitive growth assays, analyzed data, and drafted the manuscript. M.R.L. performed whole-genome sequencing and analysis and edited the manuscript. M.A.T. performed the molecular modeling and wrote the corresponding manuscript section. R.M. and S.O. provided intellectual input and project support. M.J.L., E.A.W., F.J.G., A.K.L., and D.F.W. conceived the idea of the project, reviewed the data, and drafted/edited the manuscript.

^§^Author Present Address: National Institute of Allergy and Infectious Diseases, National Institutes of Health, 9000 Rockville Pike 31 Center Dr., Bethesda, MD, 20892. The work discussed in this article was performed by R.E.K.M. elsewhere prior to becoming a federal employee. The article was written/edited by R.E.K.M. in her private capacity. No official support or endorsement by the National Institute of Allergy and Infectious Diseases is intended or should be inferred.

## References

1 Jee, Y. et al. Antimicrobial resistance: a threat to global health. The Lancet Infectious Diseases 18, 939–940, doi:https://doi.org/10.1016/S1473-3099(18)30471-7 (2018).

2 Haldar, K., Bhattacharjee, S. & Safeukui, I. Drug resistance in Plasmodium. Nat Rev Microbiol 16, 156–170, doi:10.1038/nrmicro.2017.161 (2018).

3 Phillips, M. A. & Rathod, P. K. Plasmodium dihydroorotate dehydrogenase: a promising target for novel anti-malarial chemotherapy. Infect Disord Drug Targets 10, 226–239, doi:10.2174/187152610791163336 (2010).

4 Llanos-Cuentas, A. et al. Antimalarial activity of single-dose DSM265, a novel plasmodium dihydroorotate dehydrogenase inhibitor, in patients with uncomplicated Plasmodium falciparum or Plasmodium vivax malaria infection: a proof-of-concept, open-label, phase 2a study. Lancet Infect Dis 18, 874–883, doi:10.1016/S1473-3099(18)30309-8 (2018).

5 McCarthy, J. S. et al. Safety, tolerability, pharmacokinetics, and activity of the novel long-acting antimalarial DSM265: a two-part first-in-human phase 1a/1b randomised study. Lancet Infect Dis 17, 626–635, doi:10.1016/S1473-3099(17)30171-8 (2017).

6 Murphy, S. C. et al. A Randomized Trial Evaluating the Prophylactic Activity of DSM265 Against Preerythrocytic Plasmodium falciparum Infection During Controlled Human Malarial Infection by Mosquito Bites and Direct Venous Inoculation. J Infect Dis 217, 693–702, doi:10.1093/infdis/jix613 (2018).

7 Sulyok, M. et al. DSM265 for Plasmodium falciparum chemoprophylaxis: a randomised, double blinded, phase 1 trial with controlled human malaria infection. Lancet Infect Dis 17, 636–644, doi:10.1016/S1473-3099(17)30139-1 (2017).

8 Phillips, M. A. et al. A long-duration dihydroorotate dehydrogenase inhibitor (DSM265) for prevention and treatment of malaria. Sci Transl Med 7, 296ra111–296ra111, doi:10.1126/scitranslmed.aaa6645 (2015).

9 Mandt, R. E. K. et al. In vitro selection predicts malaria parasite resistance to dihydroorotate dehydrogenase inhibitors in a mouse infection model. Sci Transl Med 11, eaav1636, doi:10.1126/scitranslmed.aav1636 (2019).

10 Phillips, M. A. et al. A long-duration dihydroorotate dehydrogenase inhibitor (DSM265) for prevention and treatment of malaria. Sci Transl Med 7, 296ra111, doi:10.1126/scitranslmed.aaa6645 (2015).

11 White, J. et al. Identification and Mechanistic Understanding of Dihydroorotate Dehydrogenase Point Mutations in Plasmodium falciparum that Confer in Vitro Resistance to the Clinical Candidate DSM265. ACS Infect Dis 5, 90–101, doi:10.1021/acsinfecdis.8b00211 (2019).

12 Hastings, I. How artemisinin-containing combination therapies slow the spread of antimalarial drug resistance. Trends Parasitol 27, 67–72, doi:10.1016/j.pt.2010.09.005 (2011).

13 Kerantzas, C. A. & Jacobs, W. R., Jr. Origins of Combination Therapy for Tuberculosis: Lessons for Future Antimicrobial Development and Application. mBio 8, doi:10.1128/mBio.01586-16 (2017).

14 Cihlar, T. & Fordyce, M. Current status and prospects of HIV treatment. Curr Opin Virol 18, 50–56, doi:10.1016/j.coviro.2016.03.004 (2016).

15 Schmid, A. et al. Monotherapy versus combination therapy for multidrug-resistant Gram-negative infections: Systematic Review and Meta-Analysis. Scientific Reports 9, 15290, doi:10.1038/s41598-019-51711-x (2019).

16 Coates, A. R. M., Hu, Y., Holt, J. & Yeh, P. Antibiotic combination therapy against resistant bacterial infections: synergy, rejuvenation and resistance reduction. Expert Review of Anti-infective Therapy 18, 5–15, doi:10.1080/14787210.2020.1705155 (2020).

17 Bodie, M. et al. Addressing the rising rates of gonorrhea and drug-resistant gonorrhea: There is no time like the present. Can Commun Dis Rep 45, 54–62, doi:10.14745/ccdr.v45i23a02 (2019).

18 Hamilton, W. L. et al. Evolution and expansion of multidrug-resistant malaria in southeast Asia: a genomic epidemiology study. Lancet Infect Dis 19, 943–951, doi:10.1016/s1473-3099(19)30392-5 (2019).

19 Manson, A. L. et al. Genomic analysis of globally diverse Mycobacterium tuberculosis strains provides insights into the emergence and spread of multidrug resistance. Nat Genet 49, 395–402, doi:10.1038/ng.3767 (2017).

20 Dheda, K. et al. The epidemiology, pathogenesis, transmission, diagnosis, and management of multidrug-resistant, extensively drug-resistant, and incurable tuberculosis. The Lancet Respiratory Medicine 5, 291–360, doi:https://doi.org/10.1016/S2213-2600(17)30079-6 (2017).

21 Gregson, J. et al. Occult HIV-1 drug resistance to thymidine analogues following failure of first-line tenofovir combined with a cytosine analogue and nevirapine or efavirenz in sub Saharan Africa: a retrospective multi-centre cohort study. Lancet Infect Dis 17, 296–304, doi:10.1016/s1473-3099(16)30469-8 (2017).

22 Hegreness, M., Shoresh, N., Damian, D., Hartl, D. & Kishony, R. Accelerated evolution of resistance in multidrug environments. Proceedings of the National Academy of Sciences 105, 13977–13981, doi:10.1073/pnas.0805965105 (2008).

23 Dean, Z., Maltas, J. & Wood, K. B. Antibiotic interactions shape short-term evolution of resistance in E. faecalis. PLoS Pathog 16, e1008278–e1008278, doi:10.1371/journal.ppat.1008278 (2020).

24 Baym, M., Stone, L. K. & Kishony, R. Multidrug evolutionary strategies to reverse antibiotic resistance. Science 351, aad3292, doi:10.1126/science.aad3292 (2016).

25 Imamovic, L. & Sommer, M. O. Use of collateral sensitivity networks to design drug cycling protocols that avoid resistance development. Sci Transl Med 5, 204ra132, doi:10.1126/scitranslmed.3006609 (2013).

26 Rosenkilde, C. E. H. et al. Collateral sensitivity constrains resistance evolution of the CTX-M-15 β-lactamase. Nat Commun 10, 618, doi:10.1038/s41467-019-08529-y (2019).

27 Maltas, J. & Wood, K. B. Pervasive and diverse collateral sensitivity profiles inform optimal strategies to limit antibiotic resistance. PLoS Biol 17, e3000515, doi:10.1371/journal.pbio.3000515 (2019).

28 Rank, G. H., Robertson, A. J. & Phillips, K. L. Modification and inheritance of pleiotropic cross resistance and collateral sensitivity in Saccharomyces cerevisiae. Genetics 80, 783–793 (1975).

29 Leroux, P., Chapeland, F., Arnold, A. & Gredt, M. New Cases of Negative Cross-resistance between Fungicides, Including Sterol Biosynthesis Inhibitors. Journal of General Plant Pathology 66, 75–81, doi:10.1007/PL00012925 (2000).

30 Dhawan, A. et al. Collateral sensitivity networks reveal evolutionary instability and novel treatment strategies in ALK mutated non-small cell lung cancer. Sci Rep 7, 1232, doi:10.1038/s41598-017-00791-8 (2017).

31 Lorendeau, D. et al. MRP1-dependent Collateral Sensitivity of Multidrug-resistant Cancer Cells: Identifying Selective Modulators Inducing Cellular Glutathione Depletion. Curr Med Chem 24, 1186–1213, doi:10.2174/0929867324666161118130238 (2017).

32 Kim, S., Lieberman, T. D. & Kishony, R. Alternating antibiotic treatments constrain evolutionary paths to multidrug resistance. Proceedings of the National Academy of Sciences 111, 14494–14499, doi:10.1073/pnas.1409800111 (2014).

33 Gonzales, P. R. et al. Synergistic, collaterally sensitive β-lactam combinations suppress resistance in MRSA. Nat Chem Biol 11, 855–861, doi:10.1038/nchembio.1911 (2015).

34 Munck, C., Gumpert, H. K., Wallin, A. I. N., Wang, H. H. & Sommer, M. O. A. Prediction of resistance development against drug combinations by collateral responses to component drugs. Science Translational Medicine 6, 262ra156–262ra156, doi:10.1126/scitranslmed.3009940 (2014).

35 Rodriguez de Evgrafov, M., Gumpert, H., Munck, C., Thomsen, T. T. & Sommer, M. O. A. Collateral Resistance and Sensitivity Modulate Evolution of High-Level Resistance to Drug Combination Treatment in Staphylococcus aureus. Molecular Biology and Evolution 32, 1175–1185, doi:10.1093/molbev/msv006 (2015).

36 Lukens, A. K. et al. Harnessing evolutionary fitness in <em>Plasmodium falciparum</em> for drug discovery and suppressing resistance. Proceedings of the National Academy of Sciences 111, 799–804, doi:10.1073/pnas.1320886110 (2014).

37 Johnson, D. J. et al. Evidence for a central role for PfCRT in conferring Plasmodium falciparum resistance to diverse antimalarial agents. Mol Cell 15, 867–877, doi:10.1016/j.molcel.2004.09.012 (2004).

38 Sisowath, C. et al. In vivo selection of Plasmodium falciparum parasites carrying the chloroquine-susceptible pfcrt K76 allele after treatment with artemether-lumefantrine in Africa. J Infect Dis 199, 750–757, doi:10.1086/596738 (2009).

39 Richards, S. N. et al. Molecular Mechanisms for Drug Hypersensitivity Induced by the Malaria Parasite’s Chloroquine Resistance Transporter. PLoS Pathog 12, e1005725, doi:10.1371/journal.ppat.1005725 (2016).

40 Cooper, R. A. et al. Alternative mutations at position 76 of the vacuolar transmembrane protein PfCRT are associated with chloroquine resistance and unique stereospecific quinine and quinidine responses in Plasmodium falciparum. Mol Pharmacol 61, 35–42, doi:10.1124/mol.61.1.35 (2002).

41 Evans, S. G. & Havlik, I. Plasmodium falciparum: effects of amantadine, an antiviral, on chloroquine-resistant and -sensitive parasites in vitro and its influence on chloroquine activity. Biochem Pharmacol 45, 1168–1170, doi:10.1016/0006-2952(93)90264-w (1993).

42 Kirkman, L. A. et al. Antimalarial proteasome inhibitor reveals collateral sensitivity from intersubunit interactions and fitness cost of resistance. Proceedings of the National Academy of Sciences 115, E6863–E6870, doi:10.1073/pnas.1806109115 (2018).

43 Stokes, B. H. et al. Covalent Plasmodium falciparum-selective proteasome inhibitors exhibit a low propensity for generating resistance in vitro and synergize with multiple antimalarial agents. PLoS Pathog 15, e1007722, doi:10.1371/journal.ppat.1007722 (2019).

44 Flannery, E. L. et al. Mutations in the P-Type Cation-Transporter ATPase 4, PfATP4, Mediate Resistance to Both Aminopyrazole and Spiroindolone Antimalarials. ACS Chemical Biology 10, 413–420, doi:10.1021/cb500616x (2015).

45 Lukens, A. K. et al. Diversity-Oriented Synthesis Probe Targets Plasmodium falciparum Cytochrome b Ubiquinone Reduction Site and Synergizes With Oxidation Site Inhibitors. J Infect Dis 211, 1097–1103, doi:10.1093/infdis/jiu565 (2014).

46 Ross, L. S. et al. In vitro resistance selections for Plasmodium falciparum dihydroorotate dehydrogenase inhibitors give mutants with multiple point mutations in the drug-binding site and altered growth. J Biol Chem 289, 17980–17995, doi:10.1074/jbc.M114.558353 (2014).

47 Ross, L. S. et al. Identification of Collateral Sensitivity to Dihydroorotate Dehydrogenase Inhibitors in Plasmodium falciparum. ACS Infectious Diseases 4, 508–515, doi:10.1021/acsinfecdis.7b00217 (2018).

48 Dong, C. K. et al. Identification and validation of tetracyclic benzothiazepines as Plasmodium falciparum cytochrome bc1 inhibitors. Chem Biol 18, 1602–1610, doi:10.1016/j.chembiol.2011.09.016 (2011).

49 Painter, H. J., Morrisey, J. M., Mather, M. W. & Vaidya, A. B. Specific role of mitochondrial electron transport in blood-stage Plasmodium falciparum. Nature 446, 88–91, doi:10.1038/nature05572 (2007).

50 Cowell, A. N. et al. Mapping the malaria parasite druggable genome by using in vitro evolution and chemogenomics. Science 359, 191–199, doi:10.1126/science.aan4472 (2018).

51 Booker, M. L. et al. Novel inhibitors of Plasmodium falciparum dihydroorotate dehydrogenase with anti-malarial activity in the mouse model. J Biol Chem 285, 33054–33064, doi:10.1074/jbc.M110.162081 (2010).

52 Guler, J. L. et al. Asexual populations of the human malaria parasite, Plasmodium falciparum, use a two-step genomic strategy to acquire accurate, beneficial DNA amplifications. PLoS Pathog 9, e1003375, doi:10.1371/journal.ppat.1003375 (2013).

53 Barbosa, C., Römhild, R., Rosenstiel, P. & Schulenburg, H. Evolutionary stability of collateral sensitivity to antibiotics in the model pathogen Pseudomonas aeruginosa. Elife 8, doi:10.7554/eLife.51481 (2019).

54 Boni, M. F., Smith, D. L. & Laxminarayan, R. Benefits of using multiple first-line therapies against malaria. Proc Natl Acad Sci U S A 105, 14216–14221, doi:10.1073/pnas.0804628105 (2008).

55 Huijben, S. & Paaijmans, K. P. Putting evolution in elimination: Winning our ongoing battle with evolving malaria mosquitoes and parasites. Evol Appl 11, 415–430, doi:10.1111/eva.12530 (2018).

56 Apjok, G. et al. Limited Evolutionary Conservation of the Phenotypic Effects of Antibiotic Resistance Mutations. Mol Biol Evol 36, 1601–1611, doi:10.1093/molbev/msz109 (2019).

57 Lozovsky, E. R. et al. Stepwise acquisition of pyrimethamine resistance in the malaria parasite. Proceedings of the National Academy of Sciences 106, 12025–12030, doi:10.1073/pnas.0905922106 (2009).

58 Dellus-Gur, E. et al. Negative Epistasis and Evolvability in TEM-1 β-Lactamase--The Thin Line between an Enzyme’s Conformational Freedom and Disorder. J Mol Biol 427, 2396–2409, doi:10.1016/j.jmb.2015.05.011 (2015).

59 Nichol, D. et al. Antibiotic collateral sensitivity is contingent on the repeatability of evolution. Nature Communications 10, 334, doi:10.1038/s41467-018-08098-6 (2019).

60 Jiao, Y. J., Baym, M., Veres, A. & Kishony, R. Population diversity jeopardizes the efficacy of antibiotic cycling. bioRxiv, 082107, doi:10.1101/082107 (2016).

61 Lambros, C. & Vanderberg, J. P. Synchronization of Plasmodium falciparum erythrocytic stages in culture. J Parasitol 65, 418–420 (1979).

62 Johnson, J. D. et al. Assessment and Continued Validation of the Malaria SYBR Green I-Based Fluorescence Assay for Use in Malaria Drug Screening. Antimicrobial Agents and Chemotherapy 51, 1926–1933, doi:10.1128/aac.01607-06 (2007).

63 Smilkstein, M., Sriwilaijaroen, N., Kelly, J. X., Wilairat, P. & Riscoe, M. Simple and Inexpensive Fluorescence-Based Technique for High-Throughput Antimalarial Drug Screening. Antimicrobial Agents and Chemotherapy 48, 1803–1806, doi:10.1128/aac.48.5.1803-1806.2004 (2004).

