## Supplemental Materials (All Supplemental Figures and Tables)) for "Collateral sensitivity as a strategy to suppress resistance emergence: the challenge of diverse evolutionary pathways"

**Supplemental Materials for ‘Collateral sensitivity as a strategy to suppress resistance emergence: the challenge of diverse evolutionary pathways’**

Data File S1: Data File S1. Individual bioreplicates of EC50 values (nM) obtained from dose response assays

Data File S2: Copy number variation analysis based on whole-genome sequencing

Data File S3: Homozygous variants identified from *in vitro* selections by whole-genome sequencing

Data File S4: Whole-genome sequencing analysis of bulk selected populations

Data File S5: Sanger sequencing of bulk TCMDC-125334 selected populations and DSM265+TCMDC-125334 selected clones

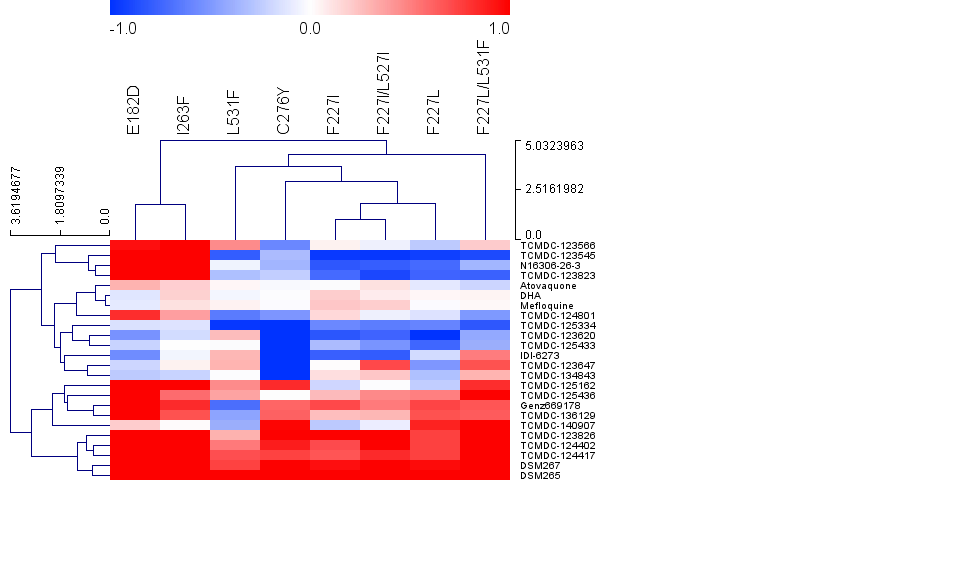

Figure S1. Hierarchical Clustering Tree of mutant lines and compounds. Clustering was done using Euclidean Distance and average linkage clustering. Data was analyzed and image produced by MultiExperimentViewer (MeV) v. 4.9.0

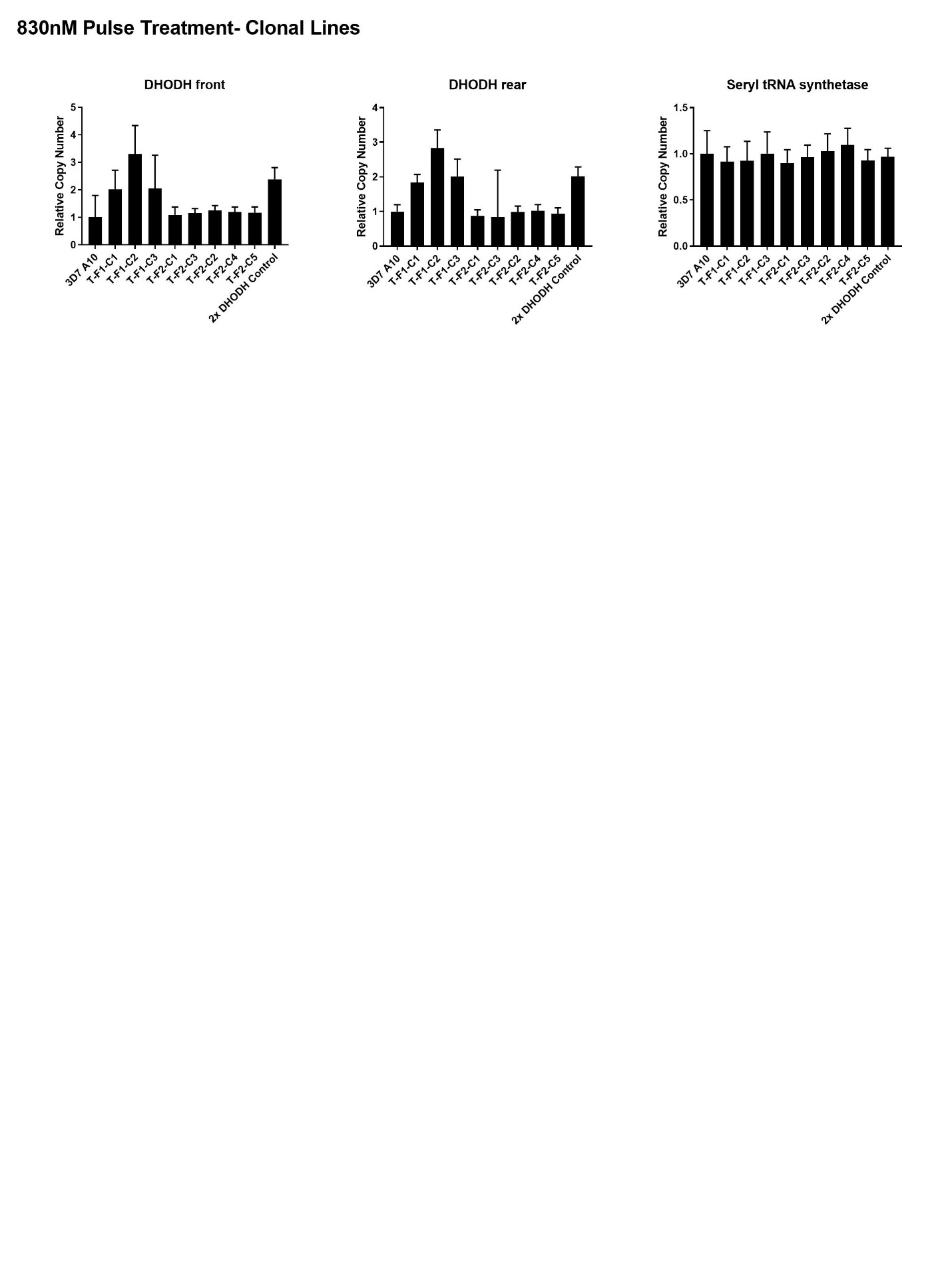

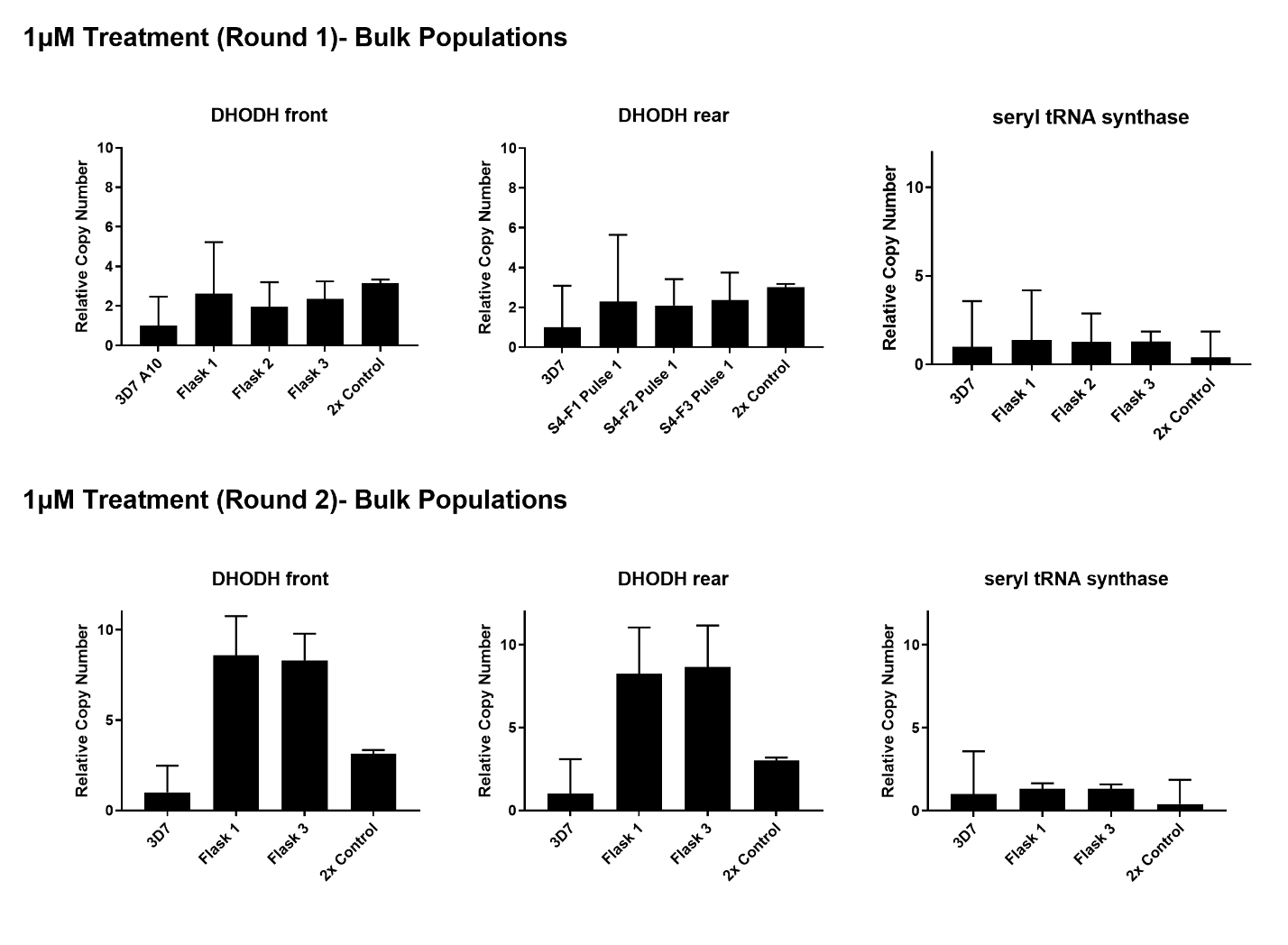

Figure S2. Copy number variation of the *dhodh* locus in 3D7 A10 parasites selected with TCMDC-125334. Copy number was detected by quantitative PCR and calculated using the ΔΔC_T_ method, normalizing to the 3D7 A10 parent and the 18s rRNA target. Seryl tRNA synthase is shown as a control. 2x DHODH is control gDNA isolated from parasites with previously-confirmed copy number duplication.

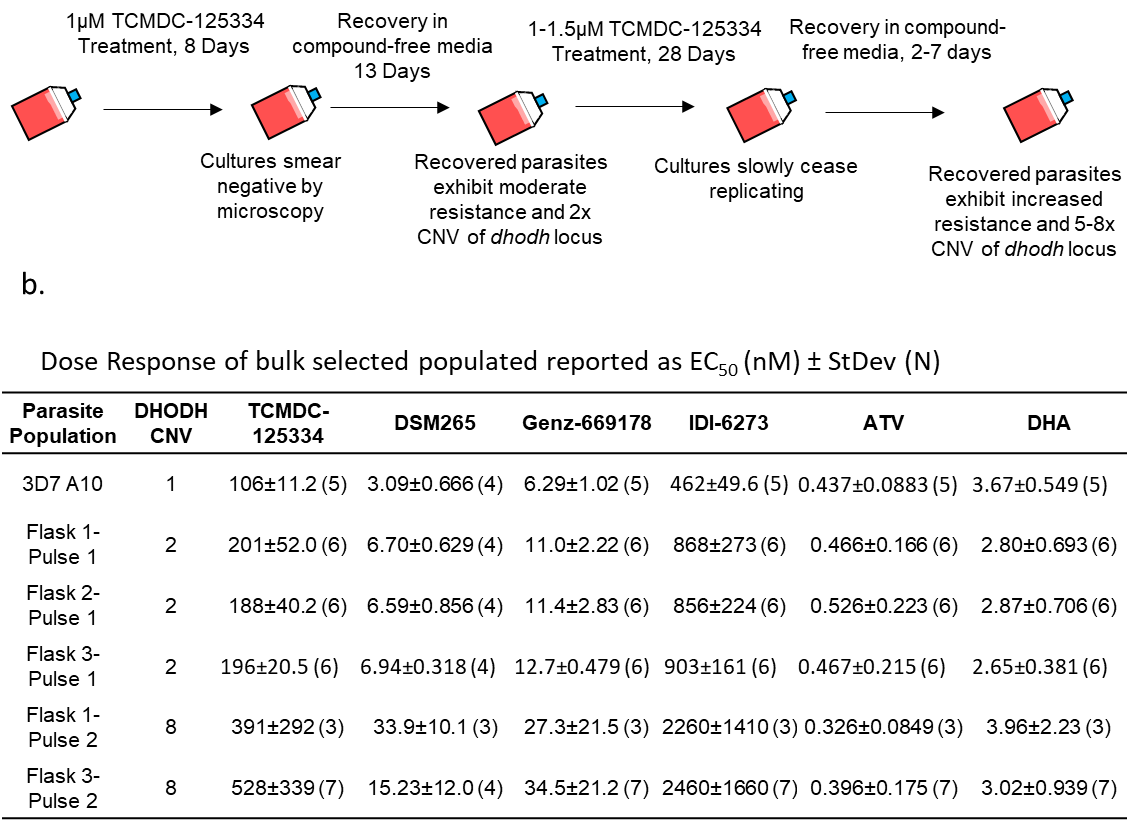

Figure S3. Treatment with 1µM TCMDC-125334 selects for moderately resistant parasite populations with copy number variation at the *dhodh* locus **A.** Schematic of selection protocol. Genetic characterization revealed that parasite populations exhibited 2x copy number variation of the *dhodh* locus after the first round of treatment with TCMDC-125334, and 5-to-8 fold copy number variation of the *dhodh* locus after the second round of treatment (see Figure S2). **B.** Dose response of parasite populations selected with TCMDC-125334. Shown are average EC_50_’s with standard deviation, and sample number (N) defined as number of individual bioreplicates.

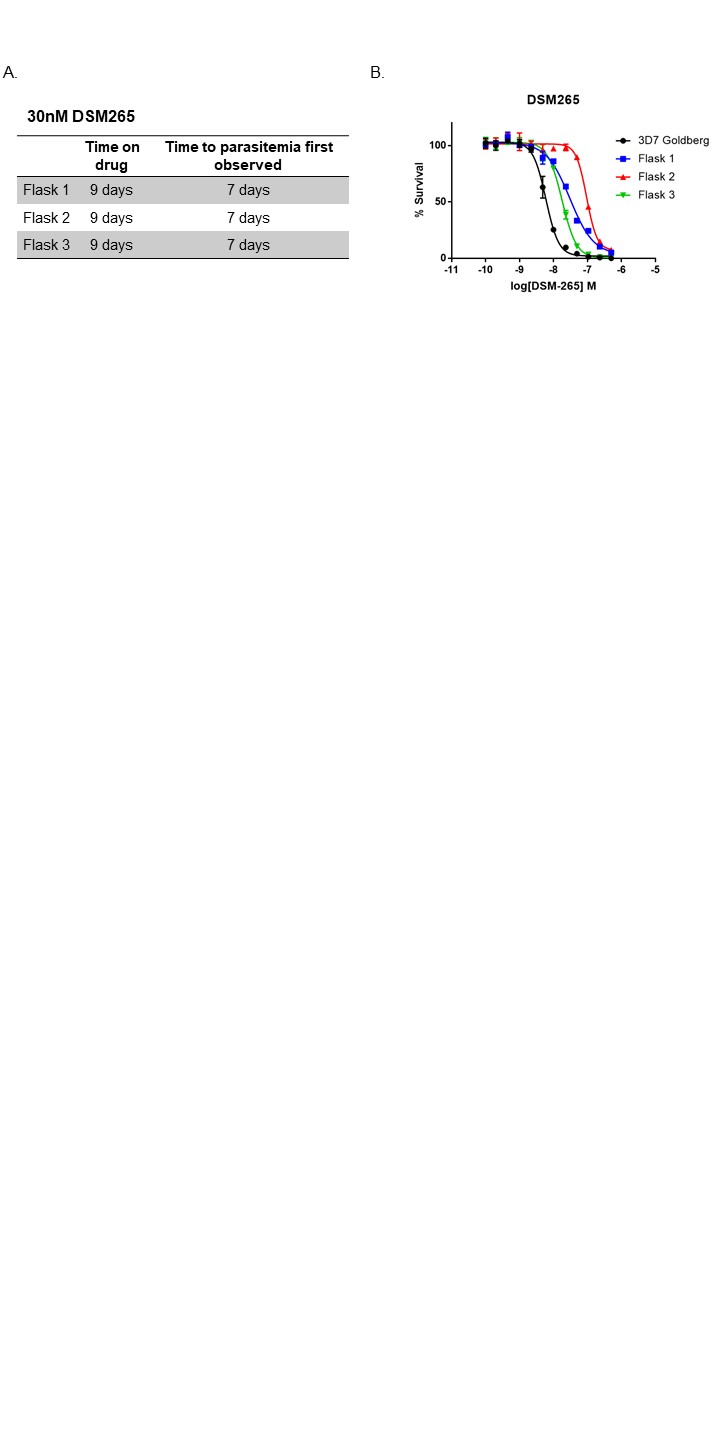

Figure S4. Control selection with DSM265. **A.** Table describing selection procedure. Three 25mL flasks of approximately 10^8^ parasites were treated with 30nM of DSM265 for nine days. In all three flasks, parasitemia was observed seven days post drug treatment. **B.** Dose response assay of *in vitro* selected bulk populations

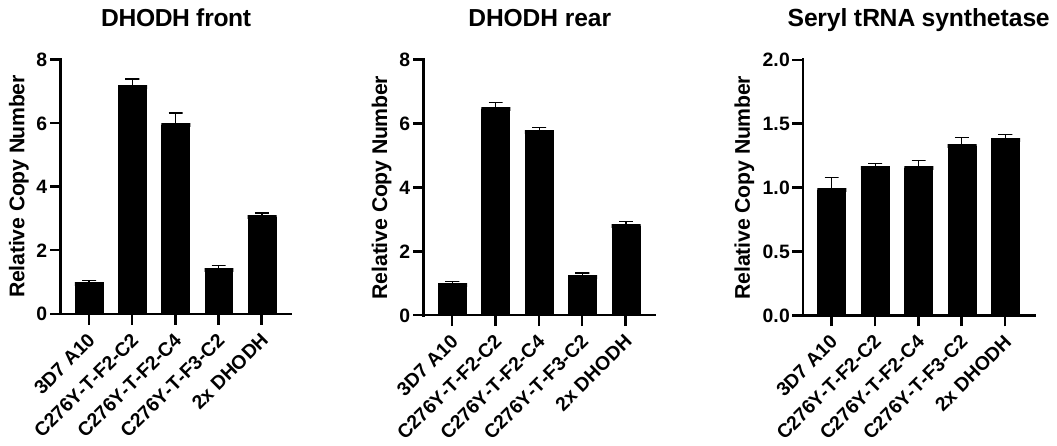

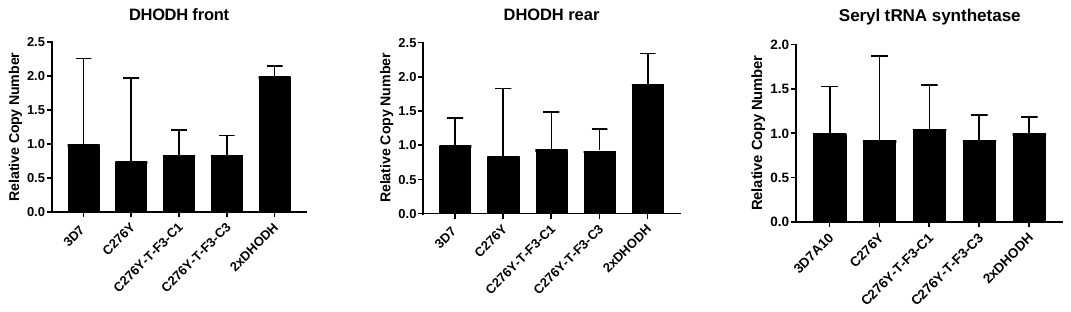

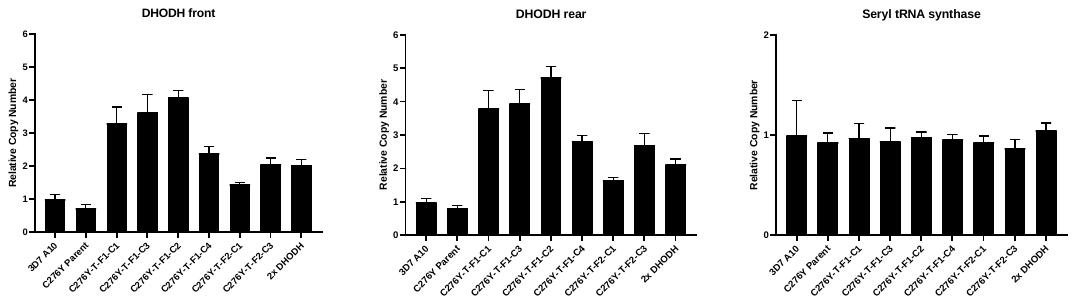

Figure S5. Copy number variation of the *dhodh* locus in DHODH C276Y parasites selected with TCMDC-125334. Copy number was detected by quantitative PCR and calculated using the ΔΔC_T_ method, normalizing to the 3D7 A10 parent and the 18s rRNA target. Seryl tRNA synthase is shown as a control. 2x DHODH is control gDNA isolated from parasites with previously-confirmed copy number duplication.

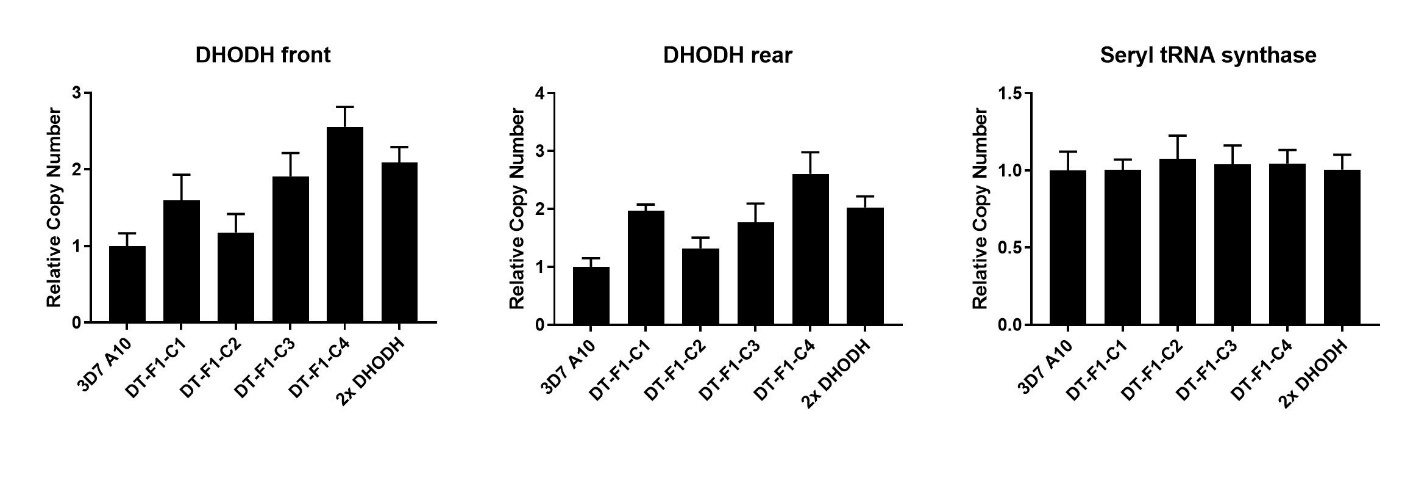

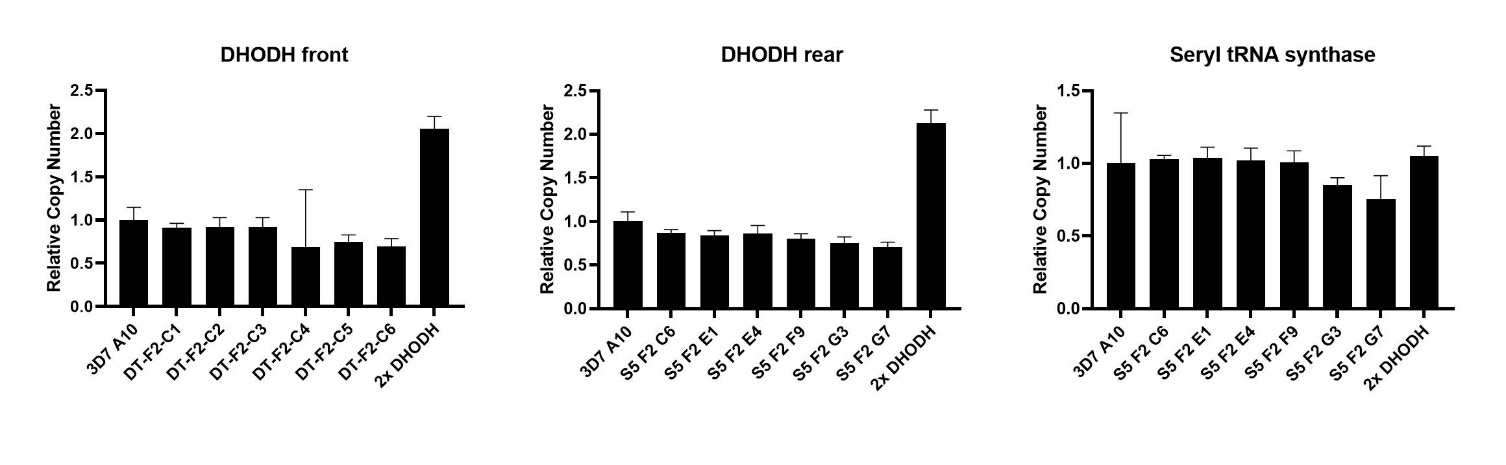

Figure S6. Copy number variation of the *dhodh* locus in 3D7 A10 parasites selected with DSM265 + TCMDC-125334 simultaneously. Copy number was detected by quantitative PCR and calculated using the ΔΔC_T_ method, normalizing to the 3D7 A10 parent and the 18s rRNA target. Seryl tRNA synthase is shown as a control. 2x DHODH is control gDNA isolated from parasites with confirmed copy number duplication.

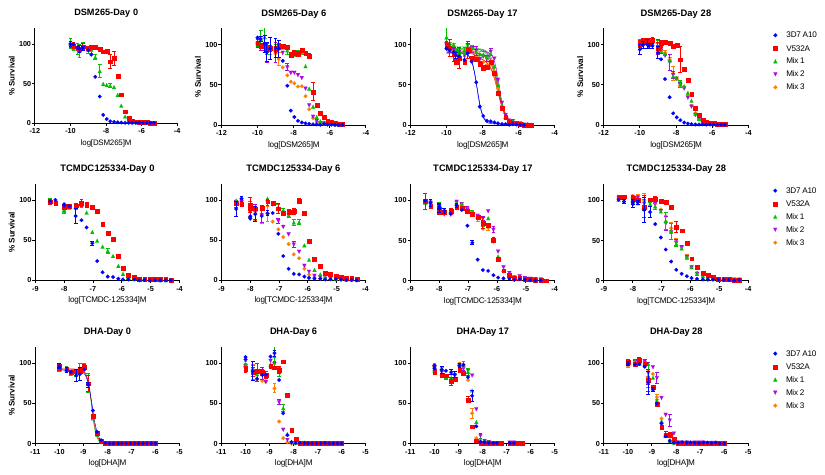

Figure S7. Co-cultured parasites in competition assay retain an intermediate phenotype. 3D7 parent and DHODH V532A mutant parasites at 1% ring stage parasitemia were mixed at equal ratio and separated into three independent flasks. Mixed cultures were grown for four weeks. Dose response assays were conducted at Day 0, Day 17 and Day 28. Upper and middle panels show the dose response for DSM265 and TCMDC-125334, respectively, which differ between the 3D7 A10 wildtype and DHODH V532A strains. The bottom panel shows the dose response for dihydroartemisinin (DHA) as a control.

**Table S1:** **DHODH mutant lines tested against Tres Cantos Antimalarial Set compounds**

| **DHODH Mutation(s)** | **Parental Line** | **Reference** |
| --- | --- | --- |
| **E182D*** | 3D7 NIH | Ross et al., 2014 |
| **F227I*** | Dd2 | Ross et al., 2014 |
| **F227I/L527I*** | Dd2 | Ross et al., 2014 |
| **F227L** | 3D7 A10 | Mandt et al., 2019 |
| **F227L/L531F** | 3D7 A10 | Mandt et al., 2019 |
| **F227Y** | 3D7^0087/N9^ | Mandt et al., 2019 |
| **I263F*** | Dd2 | Ross et al., 2014 |
| **C276Y** | 3D7 A10 | Mandt et al., 2019 |
| **L531F*** | Dd2 | Ross et al., 2014 |

*****Dose response of indicated lines were previously reported in Ross et al., 2018 [45]

**Table S2: Whole-genome sequencing statistics**

| **Sample Name** | **Aligned Reads** | **Mean Whole Genome Coverage** | **Percent Bases Covered by 5 or More Reads** |
| --- | --- | --- | --- |
| 3D7-A10-Parent | 55929898 | 170.44 | 98.8 |
| 3D7-DSM265-F1 | 54755311 | 176.06 | 98.8 |
| 3D7-DSM265-F2 | 49779616 | 158.96 | 98.6 |
| 3D7-DSM265-F3 | 46611863 | 146.6 | 98.5 |
| TCMDC125334-3D7-T-F1-C1 | 13462259 | 43.45 | 93 |
| TCMDC125334-3D7-T-F1-C2 | 39262686 | 128.49 | 98.7 |
| TCMDC125334-3D7-T-F1-C3 | 37926711 | 127.01 | 98.5 |
| TCMDC125334-3D7-T-F2-C1 | 41671469 | 134.97 | 98.5 |
| TCMDC125334-3D7-T-F2-C2 | 46419210 | 153.23 | 98.7 |
| TCMDC125334-3D7-T-F2-C3 | 49679017 | 156.84 | 98.8 |
| TCMDC125334-3D7-T-F2-C5 | 71155194 | 205.79 | 98.7 |
| TCMDC125334-3D7-T-F2-C4 | 44909308 | 145.88 | 98.6 |
| DHODH-C276Y-Parent | 68749240 | 199.72 | 98.9 |
| TCMDC125334-C276Y-T-F1-C1 | 54058057 | 168.14 | 98.6 |
| TCMDC125334-C276Y-T-F1-C3 | 49952158 | 161.41 | 98.6 |
| TCMDC125334-C276Y-T-F1-C2 | 58034702 | 182.84 | 98.7 |
| TCMDC125334-C276Y-T-F1-C4 | 63741384 | 198.26 | 98.7 |
| TCMDC125334-C276Y-T-F2-C1 | 59905480 | 189.43 | 98.7 |
| TCMDC125334-C276Y-T-F2-C3 | 50442129 | 165.79 | 98.5 |
| TCMDC125334-C276Y-T-F2-C2 | 66531226 | 205.51 | 98.8 |
| TCMDC125334-C276Y-T-F2-C4 | 71408182 | 211.07 | 98.8 |
| TCMDC125334-C276Y-T-F3-C1 | 44669436 | 148.45 | 97.8 |
| TCMDC125334-C276Y-T-F3-C2 | 77516579 | 236.97 | 98.8 |
| TCMDC125334-C276Y-T-F3-C3 | 65068538 | 206.84 | 98.7 |
| 3D7-1µM-TCMDC125334-Pulse2-F1 | 45344143 | 119.96 | 98.6 |
| 3D7-1µM-TCMDC125334-Pulse2-F3 | 58687789 | 179.22 | 98.6 |
| TCMDC125334-DSM265-3D7-DT-F1-C1 | 55428568 | 149.81 | 98.1 |
| TCMDC125334-DSM265-3D7-DT-F1-C3 | 48467511 | 134.18 | 98.6 |
| TCMDC125334-DSM265-3D7-DT-F2-C1 | 44220447 | 115.61 | 98.4 |
| TCMDC125334-DSM265-3D7-DT-F2-C2 | 54256204 | 137.56 | 98.5 |

**Table S3: Percentage of whole-genome sequencing reads calling a DHODH mutant allele in bulk populations selected with DSM265**

|  | **C276Y** | **I273M** |
| --- | --- | --- |
| Flask 1 | 5% | 14% |
| Flask 2 | 97% | 0.3% |
| Flask 3 | 0% | 44.9% |

**Table S4: Whole-genome sequencing reads of DHODH V532A allele during competitive growth experiment**

|  |  | Wildtype Reads | V532A Reads | % V532A |
| --- | --- | --- | --- | --- |
| **Flask 1** | Day 0 | 136 | 112 | 45% |
|  | Day 8 | 189 | 206 | 52% |
|  | Day 17 | 164 | 198 | 55% |
|  | Day 28 | 233 | 196 | 46% |
| **Flask 2** | Day 0 | N/A | N/A | N/A |
|  | Day 8 | 193 | 204 | 51% |
|  | Day 17 | 227 | 258 | 53% |
|  | Day 28 | 204 | 173 | 46% |
| **Flask 3** | Day 0 | N/A | N/A | N/A |
|  | Day 8 | 191 | 217 | 53% |
|  | Day 17 | 229 | 269 | 54% |
|  | Day 28 | 249 | 216 | 46% |

**Table S5: Primers used for PCR amplification and Sanger sequencing of *dhodh* locus**

| Primer Name | Primer Sequence |
| --- | --- |
| DHODH F1A | GTGTGATAGATAGCTCCAGTCG |
| DHODH R1B | CGTTTGGCCCCTTGGGGTTATGG |
| DHODH F2A | TTGATGGTGAAATATGTCATGACCTT |
| DHODH R2A | CCAAGGGCTTCTTTTTTGTTGTATTAAACC |
| DHODH F3A | GTCACATGATGAAAGATGCTAAGG |
| DHODH R3B | CGCACTTATGTGTCGCCCG |
